# Hippocampal-medial entorhinal circuit is differently organized along the dorsoventral axis in rodents

**DOI:** 10.1101/2022.05.30.493979

**Authors:** Shinya Ohara, Märt Rannap, Ken-ichiro Tsutsui, Andreas Draguhn, Alexei V. Egorov, Menno P. Witter

## Abstract

The general understanding of hippocampal circuits is that the hippocampus and the entorhinal cortex (EC) are topographically connected through parallel identical circuits along the dorsoventral axis. Our anterograde tracing and in vitro electrophysiology data, however, show a markedly different dorsoventral organization of the hippocampal projection to the medial EC (MEC). Whereas dorsal hippocampal projections are confined to the dorsal MEC and preferentially target layer Vb (LVb) over layer Va (LVa) neurons, the ventral hippocampus innervates the entire dorsoventral extent of MEC. In the ventral MEC, these projections innervate neurons in both LVa and LVb. In contrast, in the dorsal MEC, ventral hippocampal projections target mainly LVa neurons. As LVa neurons project to telencephalic structures, our findings indicate that the ventral hippocampus regulates LVa-mediated entorhinal-neocortical output from both the dorsal and ventral MEC. Overall, the marked dorsoventral differences in hippocampal-entorhinal connectivity impose important constraints on signal flow in hippocampal-neocortical circuits.

## Introduction

The entorhinal cortex (EC) constitutes a major gateway between the hippocampus and the neocortex and, together with the hippocampus, plays a crucial role in episodic memory and spatial navigation. Previous studies have shown that the hippocampus is functionally differentiated along the dorsoventral axis (Moser and Moser, 1998; Fanselow and Dong, 2010), which is accompanied by a functional gradient in EC (Strange et al., 2014). This is well characterized by the change in spatial representations of hippocampal place cells and entorhinal grid cells along the dorsoventral axis in rodents: the size of place fields increases as one moves from dorsal to ventral in the hippocampus (Jung et al., 1994; Kjelstrup et al., 2008). In parallel, the spacing between grid-cell firing locations increases from dorsal to ventral in the medial EC (MEC, Hafting et al., 2005; Brun et al., 2008; Fyhn et al., 2008). In line with such functional dorsoventral gradients, the reciprocal connections between the hippocampus and EC follow a corresponding topography: the dorsal hippocampus is connected to the dorsolateral EC, whereas the ventral hippocampus is connected to the ventromedial EC (Dolorfo and Amaral, 1998; Kloosterman et al., 2003a; Cenquizca and Swanson, 2007; Witter et al., 1989; Witter et al., 2020; van Groen et al., 2003). Despite this functional topography, the connectional organization of the entorhinal-hippocampal circuit itself is thought to be largely identical along the dorsoventral axis. Hippocampal input circuits are mediated by EC neurons in the superficial layers (layers II and III), providing inputs to all subfields of the hippocampus. In contrast, hippocampal output projections from CA1 and subiculum terminate in the deep layers of EC (layers V and VI), which in turn project to other brain regions (Witter et al., 2017).

In addition to recent reports in which the organization of the hippocampal input pathway has been further detailed (Kitamura et al., 2014; Masurkar et al., 2017), the structure of the hippocampal output circuit has been extensively revised based on the finding that the entorhinal layer V can be genetically divided into two sublayers – Va (LVa) and Vb (LVb, Ramsden et al., 2015). The two sublayers are characterized by distinct connectivity: LVa neurons project to telencephalic structures, whereas LVb neurons project locally to superficial EC layers, forming a recurrent entorhinal-hippocampal loop (Sürmeli et al., 2015; Ohara et al., 2018). It has been reported that input from the dorsal hippocampus to MEC is markedly weaker for LVa than LVb neurons and intrinsic connections from LVb to LVa neurons in the dorsal part of MEC are sparser than in the lateral EC (LEC, Ohara et al., 2021a; Rozov et al., 2020). These findings indicate that the dorsal MEC might not convey hippocampal information to the neocortex as effectively as LEC, challenging the current understanding on systems consolidation (Kitamura et al., 2017). It further raises the question what the main source of inputs to LVa neurons in the dorsal MEC might be and how MEC transfers hippocampal information to the neocortex.

In a previous study (Rozov et al., 2020), we showed that CA1 projections from the intermediate hippocampus to intermediate MEC target not only MEC-LVb but also MEC-LVa neurons. On the basis of these results we hypothesized that hippocampal innervation of MEC-LVa might differ along the dorsoventral axis, such that dorsal MEC-LVa neurons receive hippocampal input from more intermediate and/or ventral levels of the hippocampus. In this study, using anterograde tracing and in vitro electrophysiology in rodents, we experimentally tested this hypothesis by comprehensively examining the projections of CA1 and subiculum to MEC along the dorsoventral axis. We confirmed the previously reported topographical projections along the dorsoventral axis in the hippocampal-LEC circuit and showed that hippocampal outputs to LEC target both LVa and LVb regardless of the chosen dorsoventral level of origin. Importantly, we found that hippocampal projections to MEC differ profoundly along the dorsoventral axis: the dorsal hippocampus projects preferentially to dorsal MEC-LVb, whereas ventral hippocampal levels innervate MEC-LVa neurons throughout the dorsoventral extent of MEC. This organization indicates that dorsal and ventral hippocampal levels interact differently with MEC circuits along the dorsoventral axis and that the ventral hippocampus might influence the LVa-mediated cortical output of the dorsal hippocampus.

## Materials and Methods

### Animals

Projection patterns from the hippocampus to EC were anatomically characterized using adult male C57BL/6N mice (n = 6), adult male SD rats (n = 4) and adult female SD rats (n = 19). Predicted connectivity was electrophysiologically verified using brain slices obtained from 10–12 week old male C57BL/6N mice (n = 40). Mice and rats were purchased from Japan SLC (Shizuoka, Japan) or Charles River Laboratories (Sulzfeld, Germany). Animals were group housed at a 12:12 hr reversed day/night cycle and had ad libitum access to food and water. All experiments were approved either by the Center for Laboratory Animal Research, Tohoku University (Projects: 2017LsA-017; 2017LsA-018), the state government of Baden-Württemberg (Projects: G206/20; G58/21), or by the Norwegian Animal Research Authority (Projects: #8082). The experiments were conducted in accordance with the Tohoku University Guidelines for Animal Care and Use, German law, the European Communities Council Directive and the Norwegian Experiments on Animals Act.

### Surgical procedures and tracer/virus injections

For the in vivo delivery of tracers or viral vectors, animals were deeply anesthetized with vaporized isoflurane and mounted in a stereotaxic frame. Anesthesia was maintained throughout the operation by mask inhalation of isoflurane at concentrations between 1.5–2.5%. Animals used for anatomical experiments were injected subcutaneously with buprenorphine hydrochloride (0.1 mg/kg, Temgesic, Indivior), meloxicam (1 mg/kg, Metacam Boehringer Ingelheim Vetmedica), and at the incision site with bupivacaine hydrochloride (Marcain 1 mg/kg, Astra Zeneca). Mice used for electrophysiological experiments were injected with buprenorphine (0.1 mg/kg, s.c.) 30 minutes before and 3 hours after each surgery. Following head fixation, the skull was exposed and a small burr hole was drilled above the injection site. For anterograde tracing experiments, either 2.5% phaseolus vulgaris leucoagglutinin (PHA-L; Vector Laboratories, #L-1110) or 3.5–5.0% 10 kDa biotinylated dextran amine (BDA; Invitrogen, #D1956) was injected iontophoretically with positive 6–12 mA current pulses (6 s on, 6 s off) for 15 min. Alternatively, an adeno-associated virus (AAV) cocktail was used, consisting of AAV1-Syn1(S)-FLEX-tdTomato-T2A-SypEGFP (1.8 × 10^13^ GC/ml, 133 nl, Addgene #51509) and AAV9.CaMKII 0.4.Cre (2.1 × 10^13^ GC/ml, 67 nl, Addgene #105558), 200 nl of which was pressure injected using a glass micropipette (outer tip diameter = 20–40 μm) connected to a 1 μl Hamilton microsyringe. For electrophysiological experiments, 70–100 nl of AAV5-CaMKIIa-hChR2(H134R)-EYFP (UNC vector core, Karl Deisseroth virus stock/ Addgene #26969) was injected into either the ventral (AP = −2.8 – −3.1 mm; ML = ±3.4 mm; DV = −4.0 mm (21 mice)) or the dorsal hippocampus (AP = −2 mm; ML = ±2 mm; DV = −1.5 mm (10 mice) or AP = −1.5 mm; ML = ±1.2 mm; DV = −1.4 mm (9 mice)) at a rate of 200 nl per minute using a stainless steel needle (NF33BV, inner tip diameter = 115 μm) connected to a 10 μl NanoFil Syringe (WPI, Sarasota, USA). Following each injection, the pipette was left in place for another 10–15 minutes before being withdrawn. The wound was sutured and the animal was monitored for recovery from anesthesia, after which it was returned to its home cage.

### Immunohistochemistry and imaging of neuroanatomical tracing samples

Ten days after tracer or 3–4 weeks after viral injections the injected animals were anesthetized with isoflurane, euthanized with a lethal intraperitoneal injection of pentobarbital (100 mg/kg), and subsequently transcardially perfused, first with Ringer’s solution (0.85% NaCl, 0.025% KCl, 0.02% NaHCO3) and then with 4% paraformaldehyde (PFA) in 0.1 M phosphate buffer (PB). Brains were removed from the skull, post-fixed in PFA overnight, and put in a cryo-protective solution containing 20% glycerol and 2% dimethylsulfoxid (DMSO) diluted in 0.125 M PB. A freezing microtome was used to cut the brains into 40-μm-thick sections in either the coronal, horizontal, or sagittal plane, which were collected in six equally spaced series for processing.

To visualize PHA-L, sections were stained with primary (1:1000, rabbit anti-PHA-L, Vector Laboratories AS-2300) and secondary antibodies (1:400, Alexa Fluor 647 goat anti-rabbit IgG, Jackson ImmunoResearch #111-605-144), while BDA was visualized with Cy3-streptavidin (1:400, Jackson ImmunoResearch #016-160-084). GFP signal was enhanced with a primary (1:500, mouse anti-GFP, Invitrogen #A11120) and secondary antibody (1:400, Cy3 goat anti-mouse IgG, Jackson ImmunoResearch #115-165-146). For delineation purposes, sections were stained with primary (1:1000, guinea pig anti-NeuN, Millipore #ABN90P; 1:1000, mouse anti-NeuN, Millipore #MAB377; 1:300, rabbit anti-PCP4, Sigma Aldrich #HPA005792) and secondary antibodies (Alexa Fluor 647 goat anti-guinea pig IgG, Jackson ImmunoResearch #106-605-003; Alexa Fluor 647 goat anti-mouse IgG, Jackson ImmunoResearch #115-605-003; Alexa Fluor 647 goat anti-rabbit IgG).

For immunofluorescence staining, floating sections were rinsed in phosphate buffered saline (PBS) containing 0.1% Triton X-100 (PBS-Tx), followed by a 60 min incubation in blocking solution containing 5% goat serum in PBS-Tx at room temperature (RT). Sections were subsequently incubated with primary antibodies diluted in the blocking solution for 20–40 hr at 4 °C, washed in PBS-Tx (3 × 10 min), and incubated with secondary antibodies diluted in PBS-Tx for 4–6 hr at RT. Finally, sections were washed in PBS (3 × 10 min), mounted on gelatin-coated slides, and coverslipped with Entellan new (Merck Chemicals, #107961).

Sections were imaged using an automated scanner (Zeiss Axio Scan Z1). In order to precisely identify the location of the injection site in horizontally or sagittally sectioned samples, we identified the corresponding location of the injection site in the coronal plane using either the Waxholm space three-plane architectonic atlas of the rat hippocampal region (Papp et al., 2014; Boccara et al., 2015; Kjonigsen et al., 2015) or Allen Brain explorer (http://connectivity.brain-map.org/3d-viewer). To summarize the locations of the injection sites, all injection sites were transferred from the coronal plots onto an unfolded template map of CA1 and subiculum (Swanson and Hahn, 2020).

### Analysis of neuroanatomical tracing samples

The distribution of labeled axons in EC layer V was quantified either in coronal, horizontal, or sagittal sections spaced 240 μm apart. After identifying MEC and LEC and their respective layers (Ohara et al., 2018, 2021a), EC was divided into columnar bins by first dividing layer IV into 100–200 μm-wide bins and then extending the columnar bin to layers Va and Vb (Figure S1). Fluorescence intensity of immunohistochemically labeled axons within each bin was quantified using ImageJ/Fiji (Wayne Rasband, NIH, USA, open source). The intensity of immunolabeling in all bins was normalized to the bin with maximum intensity in the same sample and the normalized intensities were plotted on an unfolded map of EC layer Va/Vb. We further created a composite image to visualize the differences in labeling patterns between LVa and LVb using MATLAB (MathWorks, USA).

To compare the differences in hippocampal projection patterns between MEC-LVa, MEC-LVb, LEC-LVa, and LEC-LVb, we summed up the normalized fluorescence intensities of bins within each subregion/layer. The proportion of labeled fibers in MEC-LVa/MEC-LVb/LEC-LVa/LEC-LVb among all labeled fibers was calculated and the differences between LVa and LVb groups were tested using paired t-tests. All statistical tests were two-tailed with thresholds for significance placed at *p<0.05, **p<0.01, and ***p<0.001. All data are shown as mean ± standard errors.

### Preparation of mouse brain slices and recording of postsynaptic responses from LV neurons

Horizontal brain slices containing the hippocampus and entorhinal cortex were obtained a minimum of two weeks after virus injections using standard proceedings (Roth et al., 2016). Briefly, mice were sacrificed under deep CO2-induced anesthesia. After decapitation, brains were rapidly removed and placed in an ice-cold oxygenated cutting solution containing (in mM): 140 K-gluconate, 15 Na-gluconate, 4 NaCl, 10 HEPES, 0.2 EGTA, saturated with carbogen (95% O2 and 5% CO2, pH 7.3). 350-μm-thick horizontal slices were cut using a vibratome slicer (Leica VT1200S, Nussloch, Germany). After cutting, the slices were incubated for 20 min at ~34 °C in a lowered sodium resting solution containing (in mM): 110 NaCl, 2.5 KCl, 0.8 CaCl2, 8 MgCl2, 1.25 NaH2PO4, 26 NaHCO3, 0.4 ascorbate, 3 pyruvate, 14 glucose (pH 7.3; da Silva et al., 2019), and subsequently stored at RT in carbogen-saturated artificial cerebrospinal fluid (ACSF) containing (in mM): 124 NaCl, 3 KCl, 1.6 CaCl2, 1.8 MgSO4, 10 glucose, 1.25 NaH2PO4, 26 NaHCO3 (pH 7.4 at 34°C). Slices were allowed to recover for at least one hour before the start of electrophysiological recordings.

Individual slices were then transferred to a recording chamber which was continuously superfused with oxygenated ACSF maintained at 32 ± 1 °C. Layer Va and Vb excitatory neurons were identified with an upright microscope (BX-51 WI, Olympus, Japan) at 40× magnification using infrared-differential interference contrast (IR-DIC) microscopy. Whole-cell patch-clamp recordings were performed using borosilicate glass pipettes with a resistance of 4–6 MΩ filled with a K-based intracellular solution containing (in mM): 126 K-gluconate, 4 KCl, 10 HEPES, 0.3 EGTA, 4 Mg-ATP, 0.3 Na-GTP, 10 Na2-phosphocreatine (pH 7.3, KOH, calculated liquid junction potential −15 mV). Axonal fibers of hippocampal pyramidal cells expressing ChR2 were excited through a 40x/0.8-NA objective using a TTL-controlled blue LED (470 nm, ThorLabs no. M470L3, 1 ms pulses) and excitatory postsynaptic currents (EPSCs) were recorded in voltage-clamp mode at a holding potential of −70 mV. Data were acquired using the ELC-03XS amplifier (npi electronics, Tamm, Germany) connected to an analog-to-digital converter (POWER 1401 mkII, CED, Cambridge, UK) and stored for offline analysis using Signal4 and Spike2 (v7) software (CED, Cambridge, UK). Currents were low-pass filtered at 8 kHz and digitized at 20 kHz.

Layer Va and Vb excitatory neurons were preliminarily identified based on their location, shape of cell body and firing properties as previously described (Rozov et al., 2020). During recordings, cells were filled with biocytin (1–5%, Sigma-Aldrich, Taufkirchen, Germany, Cat. No. B4261) which was later immunolabeled to define cell morphology and location. The exact location of neurons within layer V was further verified by immunolabeling for the transcription factor Ctip2, which marks neurons in a region corresponding to sublayer Vb.

### Immunohistochemistry and imaging of recorded slices

Slices containing biocytin-filled cells were fixed in 4% PFA in PB for 45–60 min at RT and stored in PBS (pH 7.4) at 4 °C. For staining, slices were pretreated in blocking solution (5% goat serum and 0.3% Triton X-100 in PBS) for 2 hr at RT, followed by washing in PBS (3 × 15 min) and an overnight incubation (>16 hr at RT) with the primary antibody (1:1000, rat anti-Ctip2, Abcam #ab18465) diluted in antibody solution (1% goat serum and 0.2% Triton X-100 in PBS). The next day, slices were washed in PBS (3 × 15 min) and treated with the secondary antibody (streptavidin-conjugated Alexa Fluor 546 (1:1000, Invitrogen #S11225) or Alexa Fluor 647 (1:1000, anti-rat, Invitrogen #A21247)) in the antibody solution for 2 hr at RT. Slices were then washed in PBS (3 × 15 min) and incubated with 4,6-diamidino-2-phenylindole (DAPI; 1:10 000, Carl Roth, Germany) for 2 min at RT. Finally, slices were rinsed in PBS and embedded in Mowiol 4-88 (Sigma-Aldrich, Taufkirchen, Germany). Confocal image stacks were collected with a C2 Nikon confocal microscope (Nikon Imaging Center at Heidelberg University) at 2048×2048 pixel resolution (1 μm z-steps) using 4x (0.13 NA), 10x (0.45 NA) or 20x (0.75 NA) objectives in air. Multiple confocal images were merged as maximum intensity projections and analyzed with ImageJ/Fiji.

### Analysis of electrophysiological data

EPSC amplitudes were estimated as the maximum value between onset and approximate offset with subtraction of baseline level. Latencies were defined as the time interval between the onset of light pulse and the onset of EPSC. When held at −70 mV, cells with a holding current >300 pA or access resistance >40 MΩ were discarded from analysis. Because no significant differences in synaptic responses between the two dorsal hippocampal injection locations were detected, data from both groups were pooled. All data were analyzed offline using Signal4 and Spike2 (v7).

Quantitative data from multiple slices are given as median and 25th, 75th percentile [P25; P75]. Data in figures are presented as medians, [P25; P75] and individual values. Whiskers show minimum and maximum values. For values shown in figures, data in the main text are only given as medians. Statistical analysis was performed using GraphPad software (InStat, San Diego, USA). Mann-Whitney U test was used for statistical comparisons. Thresholds for significance were set identically to neuroanatomical data (ns, not significant).

## Results

### Hippocampal-entorhinal projections along the dorsoventral axis in rats

We anatomically examined the hippocampal-entorhinal projections by injecting anterograde tracers into the output structure of the hippocampal formation, either CA1 or subiculum, at different dorsoventral and proximodistal levels. The distribution of labeled fibers in EC was first examined in the rat horizontal plane (N = 14; Figure 1, S2). In line with previous studies (Naber et al., 2001; Kloosterman et al., 2003a), we observed a topographical organization of hippocampal-entorhinal projections along the proximodistal axis of the hippocampus. Anterograde tracer injections in dorsal-proximal CA1/distal subiculum resulted in labeled fibers mainly in the dorsal MEC, whereas injections in dorsal-distal CA1/proximal subiculum resulted in labeled fibers preferentially in the dorsolateral LEC. We also corroborated the laminar preference of hippocampal-MEC projections as previously reported (Sürmeli et al., 2015; Wozny et al., 2018; Ohara et al., 2018; Rozov et al., 2020), in that labeled fibers originating from dorsal-proximal CA1 mainly distributed in LVb of the dorsal MEC (Figure 1B). To examine the distribution of hippocampal-entorhinal projections in LVa and LVb, as well as along the dorsoventral and mediolateral extent of EC, we mapped the quantified label intensities on an unfolded EC map (Figure S1, see methods for detail). All samples with an injection either in dorsal-proximal CA1 or in dorsal-distal subiculum showed preferential projections to LVb rather than LVa of the dorsal MEC, and hardly any projections to LEC (Figure 1B, C, S2). This was further confirmed by quantifying the proportion of labeled fibers among all labeled fibers (Figure 1D, two-tailed paired t-test, MEC-LVa vs MEC-LVb: t4=4.895, **p=0.0081, LEC-LVa vs LEC-LVb: t4=0.4097, p=0.703). The laminar preference of hippocampal-MEC projections differed when the anterograde tracer was injected in the ventral hippocampus. In these cases, labeled axons in the ventral MEC were seen in LVa as well as in LVb, but at more dorsal levels, labeling was mainly present in LVa (Figure 1B, C, S2). As a result, the overall distribution of ventral hippocampal axons was significantly higher in MEC-LVa than in MEC-LVb, while there were no significant differences between LEC sublayers (Figure 1D, two-tailed paired t-test, MEC-LVa vs MEC-LVb: t6=7.26, **p=0.0003, LEC-LVa vs LEC-LVb: t6=1.069, p=0.3261). The ventral hippocampal projections were also characterized by their broad distribution along the dorsoventral axis of MEC, targetting not only ventral MEC but also dorsal MEC-LVa (Figure 1C, E). This is contrary to previous anatomical studies, which have reported that hippocampal-entorhinal projections are topographically organized along the dorsoventral axis: the dorsal hippocampus projects to the dorsal EC and the ventral hippocampus to the ventral EC (Kloosterman et al., 2003a; Cenquizca and Swanson, 2007).

**Figure 1.**
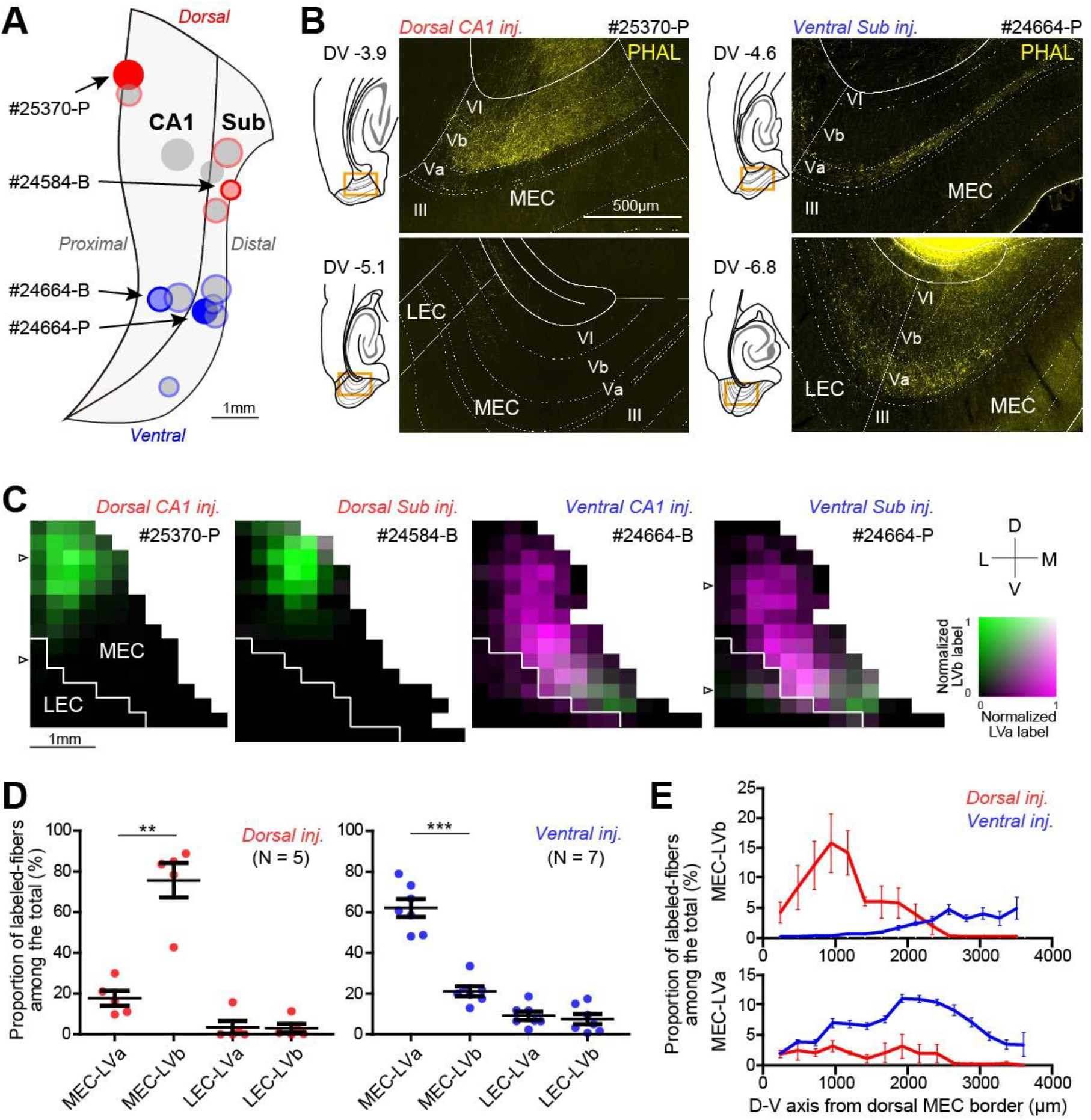
Outputs from the dorsal and ventral hippocampus target different layers and dorsoventral levels of MEC in rats. (A) Two-dimensional unfolded map of CA1 and subiculum showing the positions of anterograde tracer (PHA-L or BDA) injection sites for rat samples in the horizontal plane. Injection sites in the dorsal and ventral hippocampus are shown in red and blue, respectively. (B) Distribution of anterogradely labeled axons (yellow) in EC at different dorsoventral (DV) levels in horizontal sections. Images show samples injected either in dorsal CA1 (left, case #25370-P) or ventral subiculum (right, case #24664-P). Scale bar in the upper left panel, equalling 500 μm, holds for all four panels. (C) Four representative two-dimensional density maps showing the patterns of anterogradely labeled axons in EC following anterograde tracer injections in either dorsal or ventral CA1 or subiculum. The white arrowheads in #25370-P and 24664-P show the positions of images from B. (D) Proportion of labeled fibers among all labeled fibers in MEC-LVa, MEC-LVb, LEC-LVa, and LEC-LVb for samples injected in the dorsal (error bars: mean ± standard error; two-tailed paired t-test for MEC-LVa vs. MEC-LVb: t4 = 4.90, **p=0.0081, LEC-LVa vs. LEC-LVb: t4 = 0.41, p=0.70) and ventral hippocampus (two-tailed paired t-test for MEC-LVa vs. MEC-LVb: t6 = 7.29, ***p=0.0003, LEC-LVa vs. LEC-LVb: t6 = 1.07, p=0.33). (E) Proportion of labeled fibers in each subregion and sublayer among all labeled fibers in LVa and LVb along the dorsoventral axis of MEC.

We next examined the hippocampal projections in rat coronal sections (N = 19; Figure S3, S4) which have been widely used in previous studies to examine hippocampal-entorhinal projections (Kloosterman et al., 2003a; Cenquizca and Swanson, 2007). Samples with an injection near the border of CA1 and subiculum showed preferential projections to LEC over MEC (Figure S3A–C), similar to what we observed in horizontal sections (Figure S2). These hippocampal-LEC projections exhibited a clear topography along the dorsoventral axis: the dorsal hippocampus projected to the dorsal (lateral) LEC, and more ventrally located parts of the hippocampus projected to more ventral (medial) locations in LEC (Figure S3B–D). This topographical pattern was observed both in LEC-LVa and in LEC-LVb (Figure S3D). When the anterograde tracer was injected in either proximal CA1 or distal subiculum, labeled axons mainly distributed in MEC (Figure S4). Similar to the results in horizontal sections (Figure 1), labeled axons originating from the dorsal hippocampus mainly distributed in LVb of the dorsal (posterior) MEC. In contrast, labeled axons from the ventral hippocampus distributed in LVb in the ventral (anterior) MEC and also in LVa in the dorsal (posterior) MEC (Figure S4B, C). However, since dorsal MEC-LVa is a thin layer running parallel to the coronal plane, the projection from the ventral hippocampus to dorsal MEC-LVa was not as clear as that in the horizontal plane. Taken together, these data indicate that the previously reported topographical organization along the dorsoventral axis applies to hippocampal-LEC projections but not to hippocampal-MEC projections. In addition, the laminar preference of hippocampal-MEC projections differs between the dorsal and ventral hippocampus: the dorsal hippocampus preferentially targets MEC-LVb in the dorsal MEC, whereas the ventral hippocampus targets MEC-LVa neurons throughout the dorsoventral extent in addition to LVb in the ventral MEC.

### Hippocampal-entorhinal projections along the dorsoventral axis in mice

To examine whether the differences in hippocampal-MEC projections along the dorsoventral axis are also present in mice, we injected either BDA, PHA-L, or an AAV expressing synaptophysin-tagged-GFP into the mouse hippocampus. We then examined the resulting projection patterns in the sagittal plane (N = 9; Figure 2, S5). Similar to the results in rats, projections from the dorsal hippocampus mainly terminated in dorsal MEC-LVb among the PCP4-positive neurons, used to delineate layer Vb (Ohara et al., 2018; Figure 2B). In contrast, projections originating from the ventral hippocampus terminated mainly in the PCP4-negative MEC-LVa. Although the density of labeled axons was higher in ventral MEC-LVa than dorsal MEC-LVa, axon terminals labeled by synaptophysin-tagged-GFP were also observed in the dorsal tip of MEC (Figure 2B). This indicates that the projection from the ventral hippocampus forms synapses with LVa neurons along the entire dorsoventral extent of MEC. In addition to labeling in LVa, labeled axons also distributed in LVb in the ventral MEC (Figure 2C, S5). The overall hippocampal-MEC projection patterns, as seen in the mouse sagittal plane (Figure 2D), were consistent with those observed in rat horizontal (Figure 1), coronal (Figure S4), and sagittal samples (N = 6, Figure S6).

**Figure 2.**
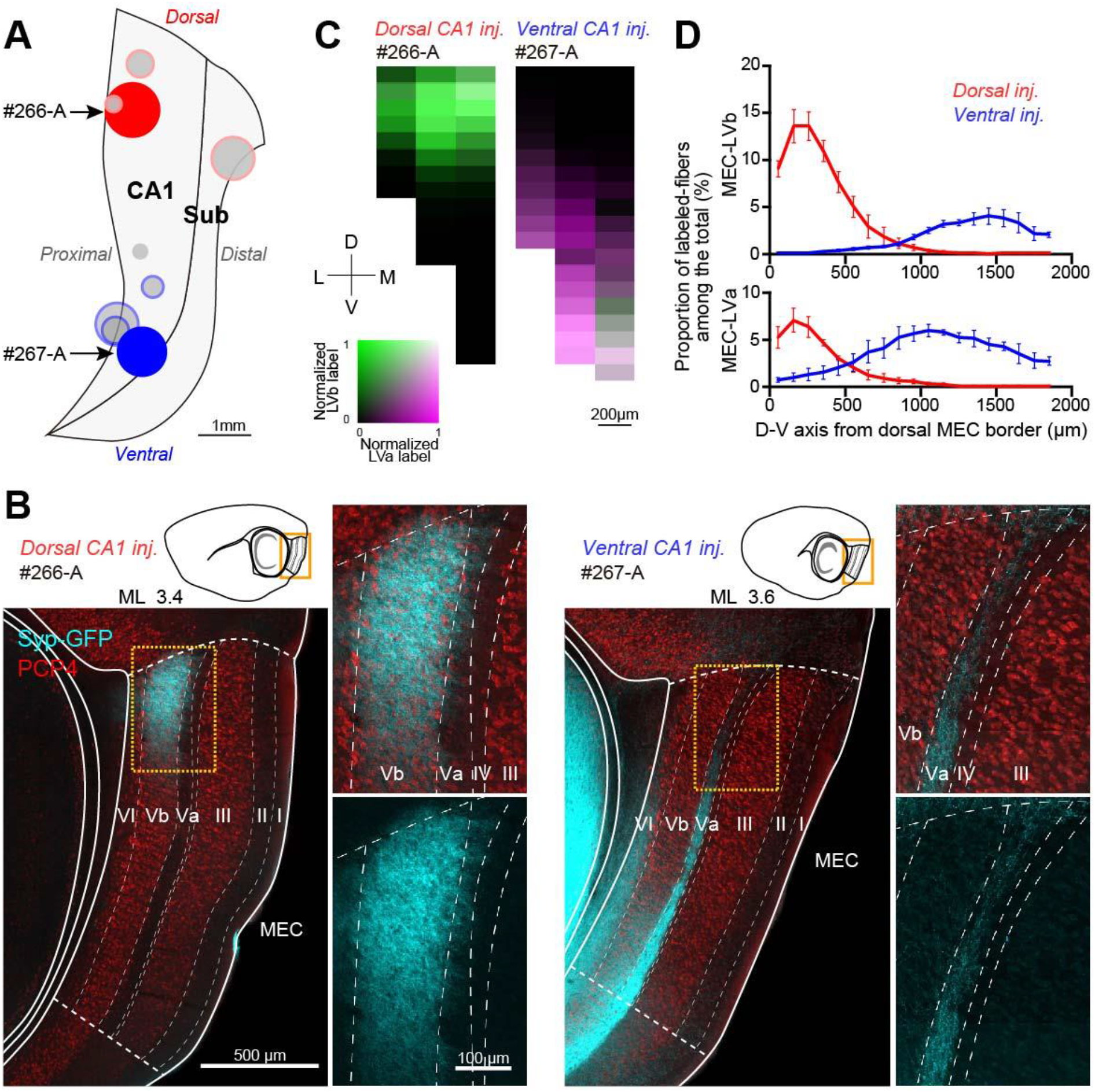
Outputs from the dorsal and ventral hippocampus target different layers and dorsoventral levels of MEC in mice. (A) Two-dimensional unfolded map of CA1 and subiculum showing the positions of anterograde tracer (PHA-L, BDA or an AAV expressing synaptophysin-tagged-GFP) injection sites in mice. (B) Fluorescent micrographs of hippocampal axons labeled with synaptophysin-tagged-GFP for samples with a dorsal (left, case #266-A) or ventral CA1 injection (right, case #267-A). Samples are immunolabeled for PCP4 to identify the PCP4-positive layers III and Vb. Scale bars in the left panel and insets also hold for the right panel and insets. (C) Two-dimensional density maps showing the patterns of anterogradely labeled axons in MEC for the two samples shown in B. (D) Proportion of labeled fibers in MEC layer Va and Vb among all labeled fibers along the dorsoventral axis of MEC.

### Ventral hippocampal outputs activate LVa neurons throughout the dorsoventral axis of MEC

As described above, anatomical tracing of hippocampal outputs to MEC suggested distinct connectivity schemes for projections from the dorsal and ventral hippocampus. Whereas dorsal hippocampal outputs preferentially target LVb and to a lesser extent LVa in the dorsal half of MEC, outputs from the ventral hippocampus form a complementary pattern, targeting LVa throughout the dorsoventral extent of MEC and LVb strongly in the ventral but not dorsal MEC. To functionally test these innervation patterns, we injected an AAV expressing hChR2-EYFP under the CaMKII promoter into either the ventral or dorsal hippocampal CA1 area. We then recorded excitatory postsynaptic currents (EPSCs) from both LVa and LVb excitatory neurons at four different levels of MEC, covering its entire dorsoventral extent, while activating ChR2-expressing hippocampal axons by illuminating LV with blue light (Figure 3A, 4A, S7A, S8A). In line with the tracing experiments, injections in ventral CA1 resulted in strong axonal fluorescence in LVa throughout all dorsoventral levels of MEC (Figure 3B, S7B). In LVb, strong fluorescent labeling was only observed at the two ventral section levels with dorsal sections displaying weak to minimal fluorescence (Figure 3B, S7B). Based on this marked discrepancy in fluorescence intensity in LVb, in our initial analysis we combined the two dorsal (section levels 3 and 4) and the two ventral sections (section levels 1 and 2) for both LVa and LVb together into a dorsal and ventral group, respectively. In the dorsal group, all 19 LVa and 16/19 LVb neurons exhibited short latency responses (LVa: 2.31 ms, n = 19; LVb: 2.45 ms, n = 16; p = 0.179, Mann-Whitney test) with comparable 20-80% EPSC rise times (LVa: 1.16 ms, n = 19; LVb: 0.91 ms, n = 16; p = 0.09, Mann-Whitney test), consistent with monosynaptic input from the ventral hippocampus. One LVb cell failed to show any discernible response, and two cells responded with longer latencies (median 5.13 ms), suggesting polysynaptic input to these cells. Importantly, short latency EPSC amplitudes were significantly larger in LVa neurons compared with LVb, reaching an over fourfold difference at higher light intensities (11.7 mW/mm2, median, LVa: −0.26 nA, n = 19; LVb: −0.06 nA, n = 16; p = 0.0001, Mann-Whitney test; Figure 3C, D left panel). To account for differences in fluorescence intensity between injections, we normalized both LVa and LVb responses to the highest LVa current amplitude at maximum light intensity in each slice. Comparison of the normalized currents revealed an even greater, roughly fivefold difference in LVa and LVb responses across most stimulation intensities (11.7 mW/mm2, median, LVa: 1.00, n = 12; LVb: 0.20, n = 13; Figure 3D, right panel). In the ventral group, all 11 LVa and 20 LVb cells responded with similarly short latencies (LVa: 2.27 ms, n = 11; LVb: 2.29 ms, n = 20; p = 0.934, Mann-Whitney test) and 20-80% EPSC rise times (LVa: 1.07 ms, n = 11; LVb: 1.02 ms, n = 20; p = 0.664, Mann-Whitney test), confirming monosynaptic innervation by the ventral hippocampus. In stark contrast to the dorsal group, however, LVb responses in ventral sections were largely comparable to LVa both in terms of absolute (11.7 mW/mm2, median, LVa: −0.41 nA, n = 11; LVb: −0.31 nA, n = 20; p = 0.223, Mann-Whitney test, Figure 3C, E left panel), as well as normalized amplitudes (11.7 mW/mm2, median, LVa: 1.00, n = 10; LVb: 0.79, n = 19; Figure 3E, right panel).

**Figure 3.**
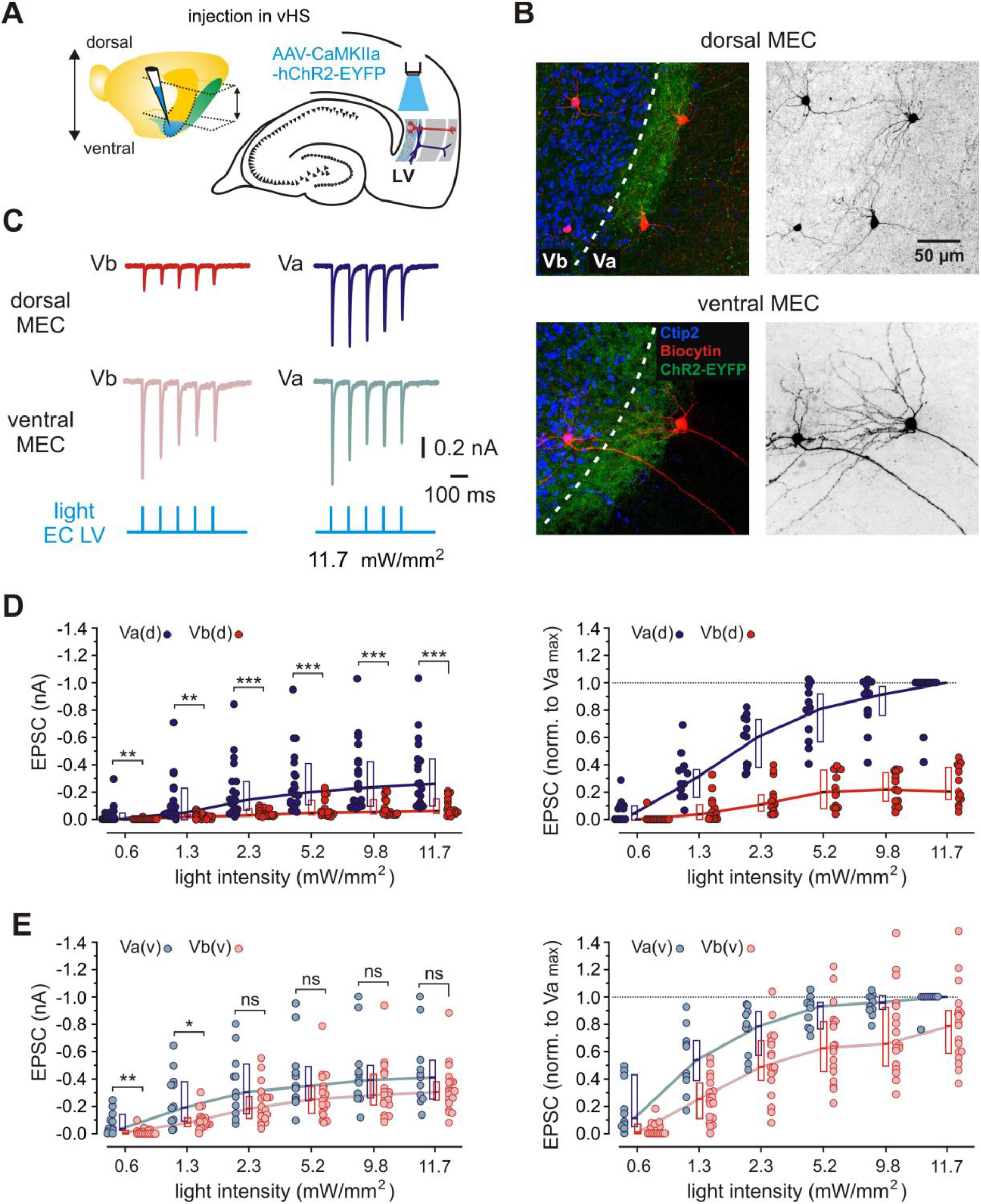
Functional connectivity between the ventral hippocampus and MEC LVa and LVb excitatory neurons. (A) Left: Illustration of the injection site (blue) in the ventral hippocampus (vHS). The approximate positions of horizontal sections used in experiments are indicated by arrows. Right: Schematic representation of a horizontal hippocampal-EC slice showing the position of light stimulation used to activate axons from ventral hippocampal neurons infected with AAV-CaMKIIa-hChR2-EYFP. (B) Z-projected confocal images of biocytin-filled MEC LVa and LVb neurons overlaid with Ctip2 immunolabeling and fluorescent staining of hippocampal axons expressing hChR2-EYFP. The dotted line indicates approximate border between LVb and LVa. Note the strong fluorescence of axonal fibers around Ctip2-negative LVa neurons in the dorsal MEC (top). Right images show the same neurons in black and white contrast. (C) Example EPSC traces recorded from LVa and LVb neurons in the same slice in the dorsal (top) and ventral (middle) MEC in response to 1 ms blue light pulses (bottom). (D) Quantification of synaptic responses of LVa and LVb neurons recorded in the dorsal MEC (LVa(d); LVb(d)). Left: plots of EPSC amplitudes induced by light pulses with increasing intensities. Right: values from the left panel normalized to the highest LVa response at maximum light intensity (11.7 mW/mm^2^) in each slice. (E) Same analysis as in (D) for LVa and LVb neurons recorded in the ventral MEC (LVa(v); LVb(v)). All data are presented as medians, *P25 and P75.* Circles represent individual values. Mann-Whitney test: \*\*\**p*<0.001; \*\**p*<0.01; \**p*<0.05; ns, not significant.

For a more detailed characterization of hippocampal input dynamics along the dorsoventral axis of MEC, we subsequently analysed LVa and LVb EPSC amplitudes at all four individual section levels (Figure S7A, B). Recordings from LVa neurons revealed strong responses at all levels of MEC with a slight but not significant trend towards smaller amplitudes in dorsal sections (11.7 mW/mm2, median, level 1: −0.41 nA, n = 5; level 2: −0.38 nA, n = 6; level 3: −0.22 nA, n = 12; level 4: −0.26 nA, n = 7; Figure S7C, top panel) (l1 vs. l2: p = 0.999; l2 vs. l3: p = 0.206; l3 vs. l4: p = 0.899; Mann-Whitney test for all). In contrast, whereas LVb neurons exhibited comparably strong responses at the two ventral section levels, EPSC amplitudes in both dorsal sections were markedly lower, reaching a sixfold difference in the dorsalmost section (11.7 mW/mm2, median, level 1: −0.31 nA, n = 11; level 2: −0.31 nA, n = 9; level 3: −0.06 nA, n = 11; level 4: −0.05 nA, n = 5; Fig. S7C, bottom panel) (l1 vs. l2: p = 0.879; l2 vs. l3: p = 0.002; l3 vs. l4: p = 0.377; Mann-Whitney test for all). Lastly, we examined LVb currents at individual section levels following normalization. As with absolute currents, there were no significant differences in amplitudes between the two ventral or the two dorsal sections but there was a sharp decline in EPSC amplitudes when transitioning from ventral to dorsal sections (11.7 mW/mm2, median, level 1: 0.87, n = 10; level 2: 0.66, n = 9; level 3: 0.23, n = 8; level 4: 0.20, n = 5; Figure S7D). (l1 vs. l2: p = 0.288; l2 vs. l3: p < 0.0001; l3 vs. l4: p = 0.724; Mann-Whitney test for all). Together, these results corroborate findings from the anatomical tracing experiments and support a connectivity scheme where ventral hippocampal outputs innervate LVa relatively uniformly throughout the dorsoventral extent of MEC but activate LVb with vastly different strength between the dorsal and ventral MEC.

### Dorsal hippocampal outputs activate LVb neurons exclusively in the dorsal MEC

We next injected AAV-hChR2-EYFP in the dorsal hippocampal CA1 area and subsequently recorded from both LVa and LVb excitatory neurons at the same four MEC levels as above (Figure 4A, S8A). In agreement with the tracing experiments and consistently with our previous report (Rozov et al. 2020), we observed strong axonal fluorescence in LVb in the dorsal half of MEC (Figure 4B, S8B). We also noted individual fibers extending towards LVa (Figure 4B, S8B) as previously described (Rozov et al. 2020). Across all injections we were unable to detect any fluorescent signal in the ventral half of MEC (Figure 4B, S8B). The lack of projections from the dorsal hippocampus to the ventral MEC was confirmed by recordings from a total of 10 LVa and 9 LVb neurons at the two ventral section levels (sections 1 and 2), all of which failed to respond to light stimulation (LVa, level 1: n = 3, level 2: n = 7; LVb, level 1: n = 3, level 2: n = 6; Figure 4C, E, S8C). In contrast, in the dorsal half of MEC (section levels 3 and 4) all 22 LVb and 26/34 LVa neurons responded to light stimulation with short latencies (LVb: 2.09 ms, n = 22; LVa: 1.96 ms, n = 26, p = 0.788, Mann-Whitney test) and similar 20-80% EPSC rise times (LVb: 0.85 ms, n = 22; LVa: 0.79 ms, n = 26; p = 0.482, Mann-Whitney test), indicating monosynaptic input from the dorsal hippocampus. Of the eight remaining LVa cells, four failed to show discernible EPSCs and four cells responded with longer latencies (median 4.99 ms), indicative of polysynaptic input. Across most light intensities, EPSC amplitudes in LVb neurons were up to threefold higher than in short latency LVa neurons (11.7 mW/mm2, median, LVb: −0.55 nA, n = 22; LVa: −0.20 nA, n = 26; p = 0.0007, Mann-Whitney test, Figure 4C, D left panel). This discrepancy in amplitudes persisted after normalizing both LVa and LVb currents to the highest LVb current amplitude at maximum light intensity in each slice, although the difference was reduced to roughly 50% across the stimulation intensities (11.7 mW/mm2, median, LVb: 1.00, n = 14; LVa: 0.48, n = 12; Figure 4D right panel). Analysis of EPSC amplitudes at the two dorsal section levels individually revealed that at both levels LVb neurons receive comparably strong input (11.7 mW/mm2, median, level 3: −0.47 nA, n = 11; level 4: −0.56 nA, n = 11; p = 0.869, Mann-Whitney test, Figure S8C bottom panel), whereas LVa responses are significantly stronger in the dorsalmost section, reaching roughly 60% of LVb amplitudes at maximum light intensity (11.7 mW/mm2, median, level 3: −0.16 nA, n = 16; level 4: −0.34 nA, n = 10; p = 0.023, Mann-Whitney test, Figure S8C top panel). This difference changed minimally after nomalization to LVb responses, with LVa EPSCs still constituting over 50% of LVb amplitudes at section level 4 (11.7 mW/mm2, median, level 3: 0.28, n = 5; level 4: 0.54, n = 7; p = 0.034, Mann-Whitney test, Figure S8D). Based on these results, dorsal hippocampal outputs exclusively innervate the dorsal half of MEC, where they exhibit a clear preference for LVb.

**Figure 4.**
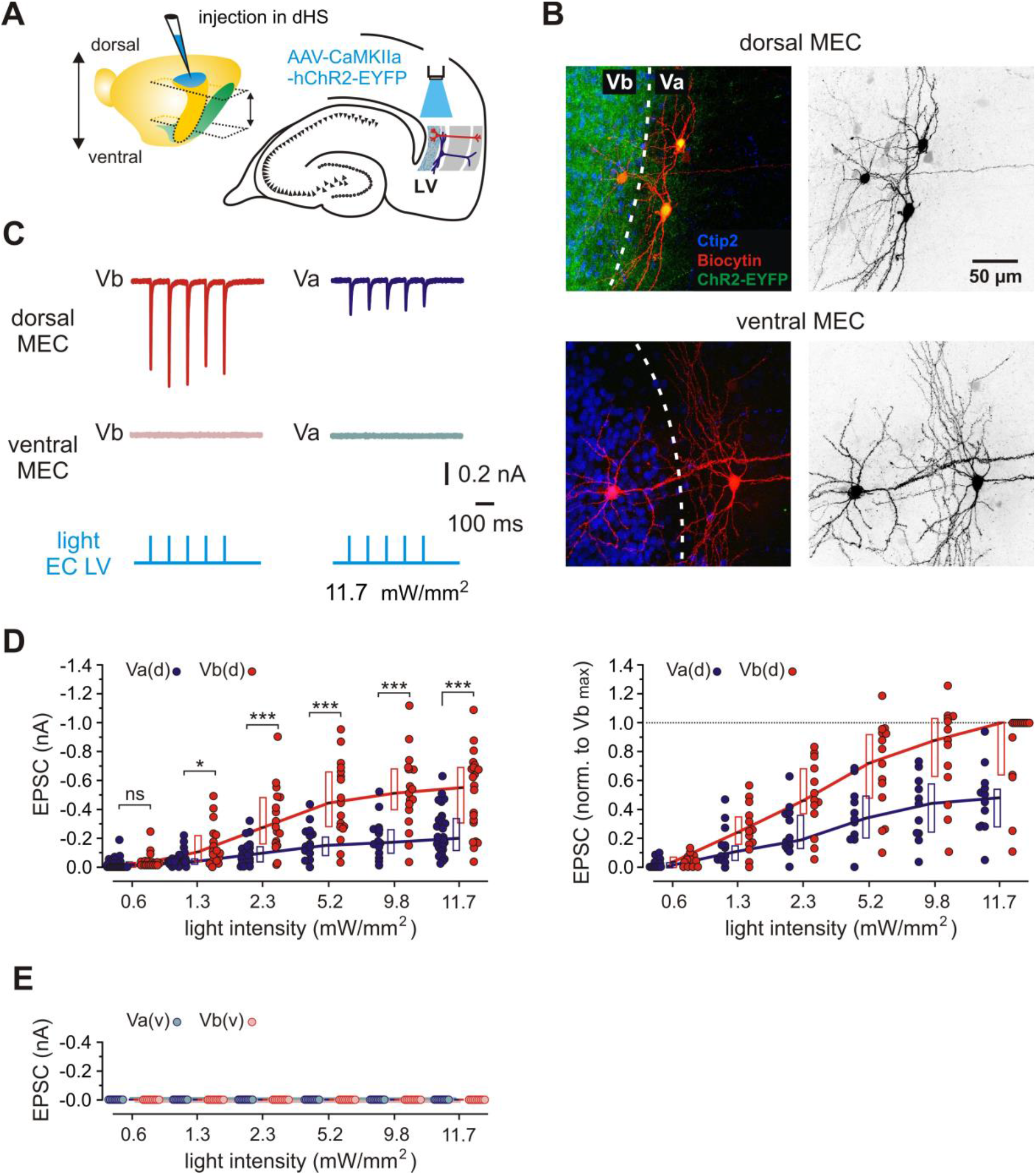
Functional connectivity between the dorsal hippocampus and MEC LVa and LVb excitatory neurons. (A) Left: Illustration of the injection site (blue) in the dorsal hippocampus (dHS). The approximate positions of horizontal sections used in experiments are indicated by arrows. Right: Schematic representation of a horizontal hippocampal-EC slice showing the position of light stimulation used to activate axons from dorsal hippocampal neurons infected with AAV-CaMKIIa-hChR2-EYFP. (B) Z-projected confocal images of biocytin-filled MEC LVa and LVb neurons overlaid with Ctip2 immunolabeling and fluorescent staining of hippocampal axons expressing hChR2-EYFP. The dotted line indicates approximate border between LVb and LVa. Note the strong fluorescence of axonal fibers in Ctip2-positive LVb and weak but recognizable fluorescence around the Ctip2-negative LVa neuron in the dorsal MEC (top), and no fluorescence in the ventral MEC (bottom). Right images show the same neurons in black and white contrast. (C) Example EPSC traces recorded from LVa and LVb neurons in the same slice in the dorsal (top) and ventral (middle) MEC in response to 1 ms blue light pulses (bottom). (D) Quantification of synaptic responses of LVa and LVb neurons recorded in the dorsal MEC (LVa(d); LVb(d)). Left: plots of EPSC amplitudes induced by light pulses with increasing intensities. Right: values from the left panel normalized to the highest LVa response at maximum light intensity (11.7 mW/mm^2^) in each slice. (E) Same analysis as in (D) for LVa and LVb neurons recorded in the ventral MEC (LVa(v); LVb(v)). All data are presented as medians, *P25 and P75.* Circles represent individual values. Mann-Whitney test: \*\*\**p*<0.001; \**p*<0.05; ns, not significant.

## Discussion

Our data reveal major differences in hippocampal-entorhinal connectivity along the dorsoventral axis of the rodent brain. By combining anatomical and electrophysiological approaches we found new, highly specific projection patterns from different portions of the hippocampus to both MEC and LEC. While projections to LEC maintain a strict longitudinal organization, projections to MEC exhibit a hitherto unknown widespread connectivity pattern. Two findings are particularly important: (1) Hippocampal projections to MEC are not restricted to a single dorsoventral level; most prominently, the ventral hippocampus targets both the ventral and dorsal MEC. (2) The specific targets of hippocampal-entorhinal projections differ along the dorsoventral axis: whereas dorsal hippocampal projections excite mainly layer Vb neurons in the dorsal MEC, ventral projections excite LVa neurons in both the ventral and dorsal MEC. This architecture has strong functional implications for the transfer of information between hippocampal networks and the neocortex.

The output of patterned activity from the hippocampus to the deep layers of EC constitutes a major pathway for the transfer of information to associative neocortical networks, which likely supports the consolidation of transiently stored information into long-term memory (Siapas and Wilson, 1998; Girardeau et al., 2009; Khodagholy et al., 2017). At the same time, the activity patterns also propagate to superficial EC layers from where they may return to the hippocampus, forming a recurrent feedback loop (Iijima et al., 1996; Kloosterman et al., 2003b; Yamamoto and Tonegawa, 2017). These two parallel pathways are associated with two different cell populations in the deep EC, namely LVa, which harbors projection neurons targeting further neocortical areas, and LVb, containing projection neurons targeting superficial EC layers (Ohara et al., 2018, 2021b).

How the hippocampus projects to these two cell populations has remained controversial. A pioneering study on hippocampal outputs to MEC reported that projections from hippocampal CA1 selectively target LVb neurons (Sürmeli et al., 2015). Subsequent work showed that both LVa and LVb neurons are targeted but the strength of innervation depends on the longitudinal position of the hippocampal origin (Rozov et al., 2020). The present study provides a comprehensive account of the three-dimensional structure of the hippocampal-entorhinal output pathway. We show that neurons from ventral hippocampal levels project to both LVa and LVb in the ventral MEC but have a strong preference for LVa neurons in the dorsal MEC (Figure 5A blue). Neurons from dorsal hippocampal portions, in contrast, project exclusively to the dorsal MEC where they exhibit a strong preference for LVb over LVa (Figure 5A red).

**Figure 5.**
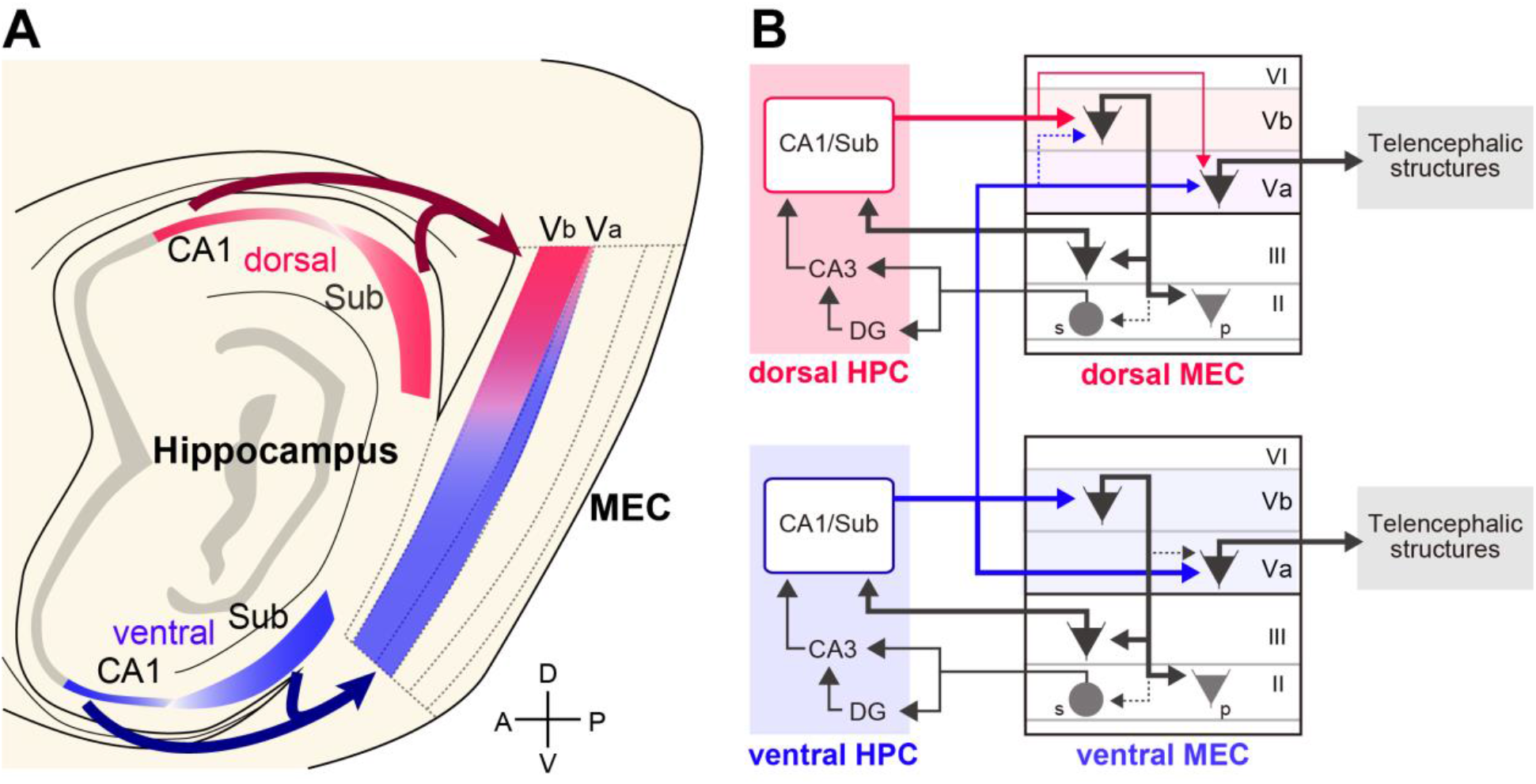
Schematic diagram of hippocampal output circuits via the medial entorhinal cortex (MEC). (A) Dorsal proximal CA1/distal subiculum innervate both layer Va and Vb neurons in the dorsal MEC with a marked preference for layer Vb neurons. Ventral proximal CA1/distal subiculum, in turn, innervate both layer Va and Vb neurons in the ventral MEC and preferentially layer Va neurons in the dorsal MEC. (B) Information from the dorsal hippocampus is mainly sent back to the hippocampus through the MEC layer Vb → MEC layer III → hippocampus loop circuit. In contrast, information from the ventral hippocampus is sent out to telencephalic structures via the MEC layer Va output circuit. Note that dorsal hippocampal information can also reach telencephalic structures through layer Va in the dorsal MEC. s: stellate cell; p: pyramidal cell.

We found projections from ventral levels of the hippocampus to the dorsal MEC, contradicting previous anatomical studies which reported a strict topographic organization along the dorsoventral axis where the dorsal hippocampus projects to dorsolateral and the ventral hippocampus to ventromedial portions of EC (Kloosterman et al., 2003a). While our present data confirm this topographical pattern for projections to LEC, they show a much more widespread projection from the ventral hippocampus to MEC. Indeed, outputs from the ventral hippocampus reach the entire MEC, including its most dorsal portion. The inconsistency between our present and previous findings is likely caused by technical differences. Most previous studies examined hippocampal-entorhinal connectivity in coronally cut sections, which are unsuited for accurately evaluating labeling patterns in the dorsal MEC. This is especially the case for LVa which in the dorsal MEC is particularly thin and can be easily overlooked in the coronal plane. Of note, one previous study did describe a dense band of terminal ventral CA1 fibers along the dorsoventral extent of MEC. This band, located in layers

IV and V of the deep MEC in the nomenclature used by the authors, likely corresponds to our layer Va (Cenquizca and Swanson, 2007, Figure 10D and 11B). These data are thus in line with our present findings, even though the authors supported a topographical distribution of CA1 projections as described above.

At a larger scale, the innervation of dorsal MEC LVa neurons by ventral hippocampal inputs is at odds with the long-postulated separation between the dorsal two-thirds and the ventral one-third of the hippocampus (Moser and Moser, 1998). This notion was based on the fact that associational fibers of dentate hilus (Fricke and Cowan, 1978) and CA3 (Swanson et al., 1978) tend to stay within the limits of their respective dorsal-ventral portions of the hippocampal formation (for a review see Witter et al., 1989). Our present findings, however, point to a convergence of projections from the dorsal and ventral hippocampus in the dorsal MEC, allowing for the integration of this apparently segregated information before being projected to downstream telencephalic networks via LVa neurons. Whether the anatomical segregation between inputs to LVa and LVb neurons is translated into a functional segregation depends on the cross-talk between the populations.

In our previous studies, we have shown that LVb neurons in the dorsal MEC strongly innervate CA1-projecting layer III neurons but have only sparse connections to LVa (Rozov et al., 2020; Ohara et al., 2018, 2021a). These findings, together with the present results, suggest that the dorsal hippocampus primarily activates the LVb-mediated hippocampal-MEC-hippocampal loop, supporting reverberating activity patterns (Figure 5B top). In contrast, the ventral hippocampus targets the hippocampal output circuit by innervating LVa neurons throughout all dorsoventral levels of MEC, which in turn send projections to various telencephalic structures (Figure 5B bottom). A part of this signal, however, could be transmitted back to the hippocampus as MEC LVa neurons were recently shown to target pyramidal neurons in the hippocampal CA1 region (Tsoi et al., 2021).

The dorsal hippocampus is strongly involved in cognitive processes such as spatial, episodic and declarative memory formation, requiring interactions with downstream neocortical areas (Fanselow and Dong, 2010). How can this be achieved within the connectivity scheme outlined above? There are several possible routes for the transfer of information from the dorsal hippocampus to telencephalic structures. First, activity might propagate from dorsal to ventral hippocampal levels through intrinsic circuits. Indeed, there is a well-developed, longitudinally projecting synaptic network among CA1 pyramidal neurons (Amaral and Witter 1989; Yang et al., 2014), from where activity can be directly routed to the medial prefrontal cortex through strong projections emerging from neurons in CA1 and subiculum (Jay and Witter, 1991; Hoover and Vertes, 2007). In addition, neurons in the dorsal subiculum send strong excitatory projections to the retrosplenial cortex (Witter et al., 1990; Bienkowski et al., 2018; Cembrowski et al., 2018), complemented by weaker long-range GABA-ergic projections from dorsal levels of CA1 (Miyashita and Rockland, 2007).

Alternatively, and at the core of the present study, hippocampal output is fed into the entire EC LVa output circuit. In LEC, this output pathway is strictly topographically organized such that the dorsoventral axis of the hippocampus is precisely mapped onto the dorsoventral axis of LEC, enabling direct access to the telencephalon. In MEC, the LVa-mediated telencephalic pathway comprises two components. First, LVa neurons in the dorsal MEC receive weak monosynaptic input from the dorsal hippocampus, in line with our previous report (Rozov et al., 2020) and in contrast to previous studies (Sürmeli et al., 2015 Wozny et al., 2018). Although these inputs are markedly weaker for LVa than LVb neurons, their presence opens the possibility for direct outputs from the dorsal hippocampus to the neocortex. Secondly, the present study shows that LVa neurons in the dorsal MEC receive additional inputs from the ventral hippocampus. The weaker activation of LVa in the dorsal MEC, roughly 50% of that seen in LVb, could potentially be compensated by projections from the ventral hippocampus as stimulation of ventral hippocampal terminals in the dorsal MEC quantitatively matched the “missing” ~ 50% of excitation in LVa. Synchronous input from both the dorsal and ventral hippocampus to dorsal MEC LVa would thus equal the strength of excitation from the dorsal hippocampus to LVb, rendering the output to telencephalic or superficial entorhinal networks similarly efficient. It is currently unknown how many dorsal LVa neurons are simultaneously innervated by both dorsal and ventral parts of the hippocampus. In our electrophysiological recordings, 76% of tested LVa neurons showed monosynaptic responses to dorsal hippocampal input and all tested LVa neurons showed monosynaptic responses to ventral input. It is therefore highly likely that a substantial fraction of dorsal MEC LVa neurons integrates dorsal and ventral hippocampal signals.

What might be the functional relevance of the convergent hippocampal afferents in the dorsal MEC? Multiple lines of evidence show that the ventral hippocampus processes information related to emotion, stress, and motivation and is a critical hub for networks that process emotion-related learning (Fanselow and Dong, 2010). These memory-related processes are supported by different state-dependent types of network oscillations, namely sharp wave-ripple complexes (SPW-Rs) for the output of hippocampal activity patterns to downstream areas (Buzsáki, 1986; 2015), and theta oscillations for coordination of network activity during acquisition of information (O’Keefe, 1993). Both patterns can propagate throughout the dorsoventral axis of the hippocampus (Patel et al., 2012; Patel et al., 2013; Lubenov and Siapas, 2009), supporting the integration of information from both the dorsal and ventral portion. Maurer and Nadel (2020) recently proposed that communication along the dorsoventral hippocampal axis is instrumental for the network to be able to recognize changes in context, whereby the ventral hippocampal output would signal such contextual change. The projection from the ventral hippocampus to the dorsal MEC, reported in our present study, would thus allow the dorsal MEC LVa neocortical output pathway to contain both contextual and salience information. This mechanism may underlie the recent finding that the projection from MEC LVa to the medial prefrontal cortex is essential for consolidating contextual fear memories (Kitamura et al., 2017). Our anatomical data indicate that a similar ventral-hippocampus-dependent transfer of hippocampal output is unlikely to be present in the LEC LVa output pathway. It is important to note, however, that according to a recent report SPW-Rs may also occur asynchronously in the dorsal and ventral hippocampus of rats (Sosa et al., 2020). Hippocampal activity could thus also be read out independently from the ventral or dorsal portion, supporting selective activation of emotion-related versus neutral downstream processes.

In summary, our findings reveal differential, yet partially convergent pathways for dorsal and ventral hippocampal outputs to the entorhinal cortex. The different cognitive, behavioral and emotional functions of dorsal and ventral hippocampal portions are reflected in distinct connectivity to downstream networks which are likely to excite the hippocampal feedback loop in some situations, and feedforward telencephalic projections in others. At the same time, there is a complex interplay between both pathways, supported by convergent connections and multiple excitatory loops at different levels. As a result, the complex nested hippocampal-entorhinal output network may be instrumental in the integration of emotional, spatial, contextual, as well as episodic contents. Together, these inputs may provide conditional gating of information to neocortical networks mediating long-term memory formation and storage.

## Acknowledgements

This work was supported by the following grants and organization: Grants-in-Aid for Scientific Research on Innovative Areas (21H00178 to S.O.), for Scientific Research (19K06917 to S.O.) from the Japan Society for Promotion of Science, PRESTO (JPMJPR21S3 to S.O.) from the Japan Science and Technology Agency, the Deutsche Forschungsgemeinschaft (DFG, German Research Foundation) – No. 430282670 (EG134/2-1) to A.V.E. Further support (to MPW) was provided by The Kavli Foundation and the Norwegian Research Council: infrastructure grant NORBRAIN #197467 and the Centre of Excellence scheme – Centre for Neural Computation #223262.

## Authors Contributions

S.O. collected and analyzed the anatomical data. M.R. carried out whole-cell recordings. M.R. and A.V.E. performed the electrophysiological and corresponding morphological data analysis. All authors conceptualized the study, interpreted the results, and wrote the manuscript.

