## Supplementary figures for "Hippocampal-medial entorhinal circuit is differently organized along the dorsoventral axis in rodents"

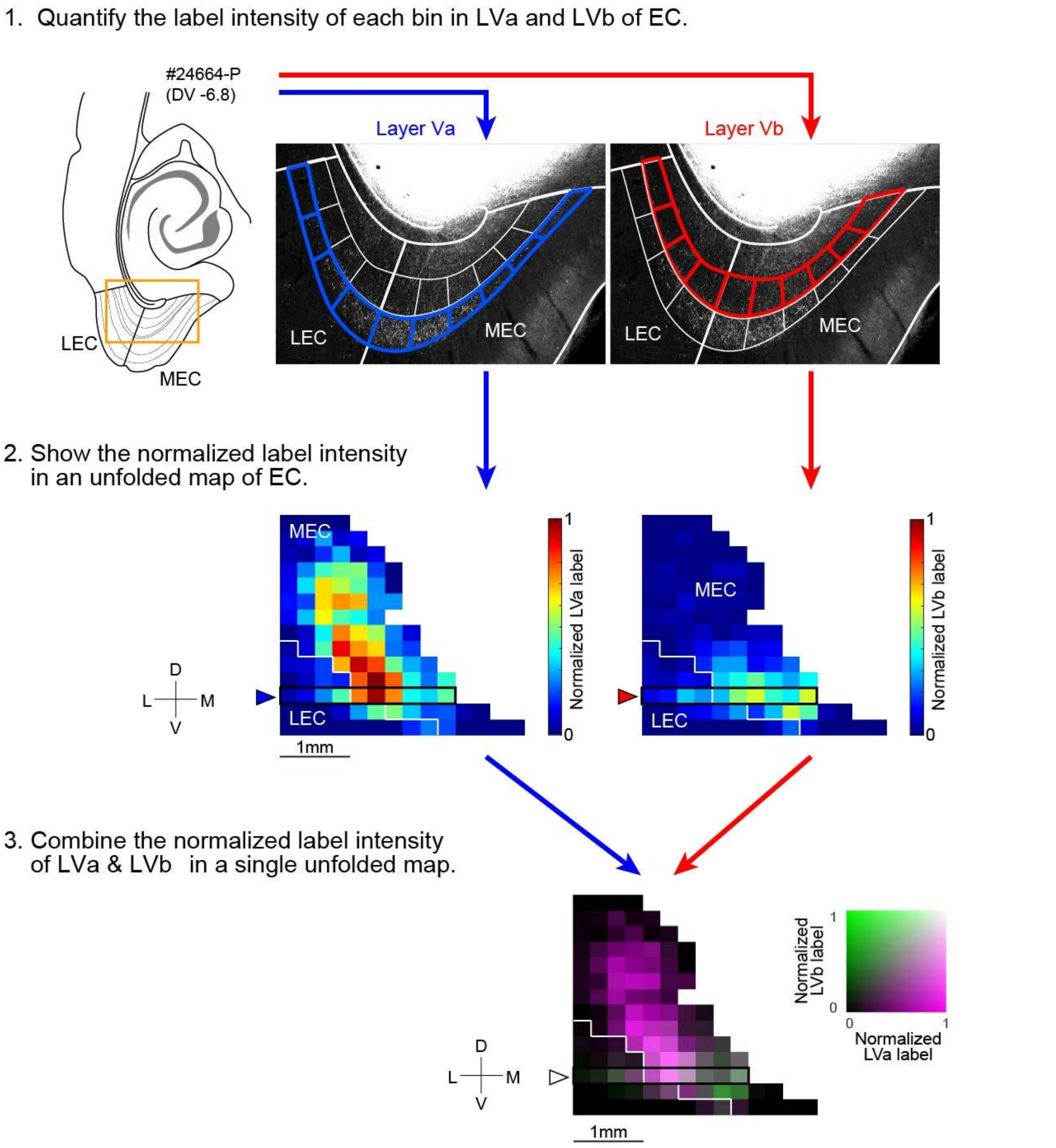


Figure S1. Schematic diagram illustrating the quantitative analysis of labeled axons in EC LVa and LVb. Samples were analyzed in sections spaced 240 μm apart in either the coronal, sagittal, or horizontal plane. Layers Va and Vb were divided into columnar bins and the label intensity of each bin was quantified (step 1). The intensity of each bin was then normalized for every sample and the normalized intensities were mapped on an unfolded map of EC (step 2). Lastly, normalized label intensities for layers Va and Vb were combined in a single unfolded map (step 3). Green indicates bins with a dense labeling of axons in LVb, while Magenta indicates bins with densely labeled axons in LVa. Bins with a dense labeling of axons in both LVa and LVb are shown in white, while bins with no labeled axons are shown in black.

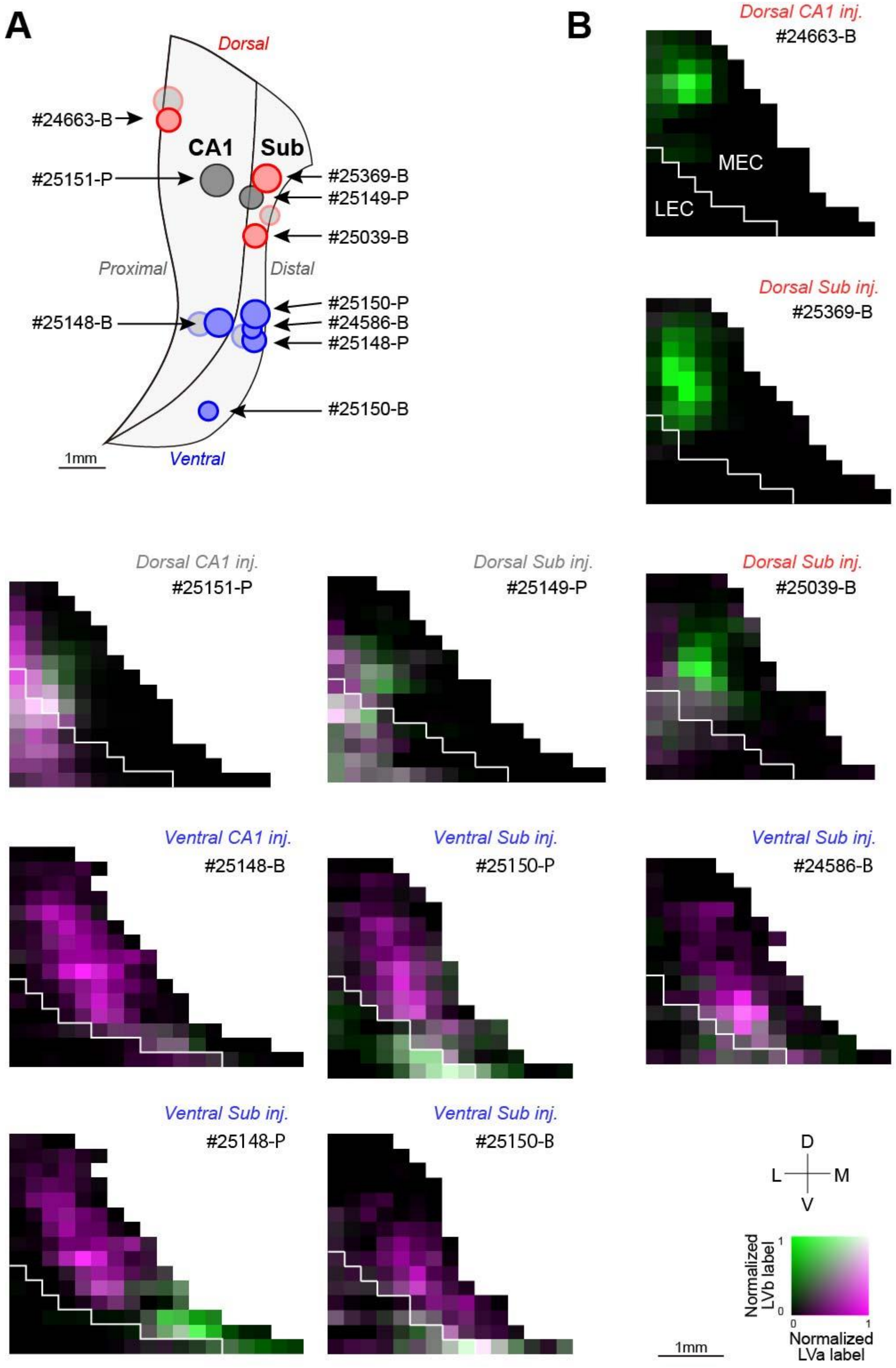


Figure S2. Related to Figure 1, shows label distribution patterns in unfolded EC maps for rat horizontal samples. Two samples, one with an injection in dorsal distal CA1 (#25151-P) and the other in dorsal proximal CA1 (#25149-P), showed massive labeling of axons in LEC. Due to this labeling pattern, these two samples were excluded from the analysis in Figure 1D, E which focuses on labeling patterns in MEC.

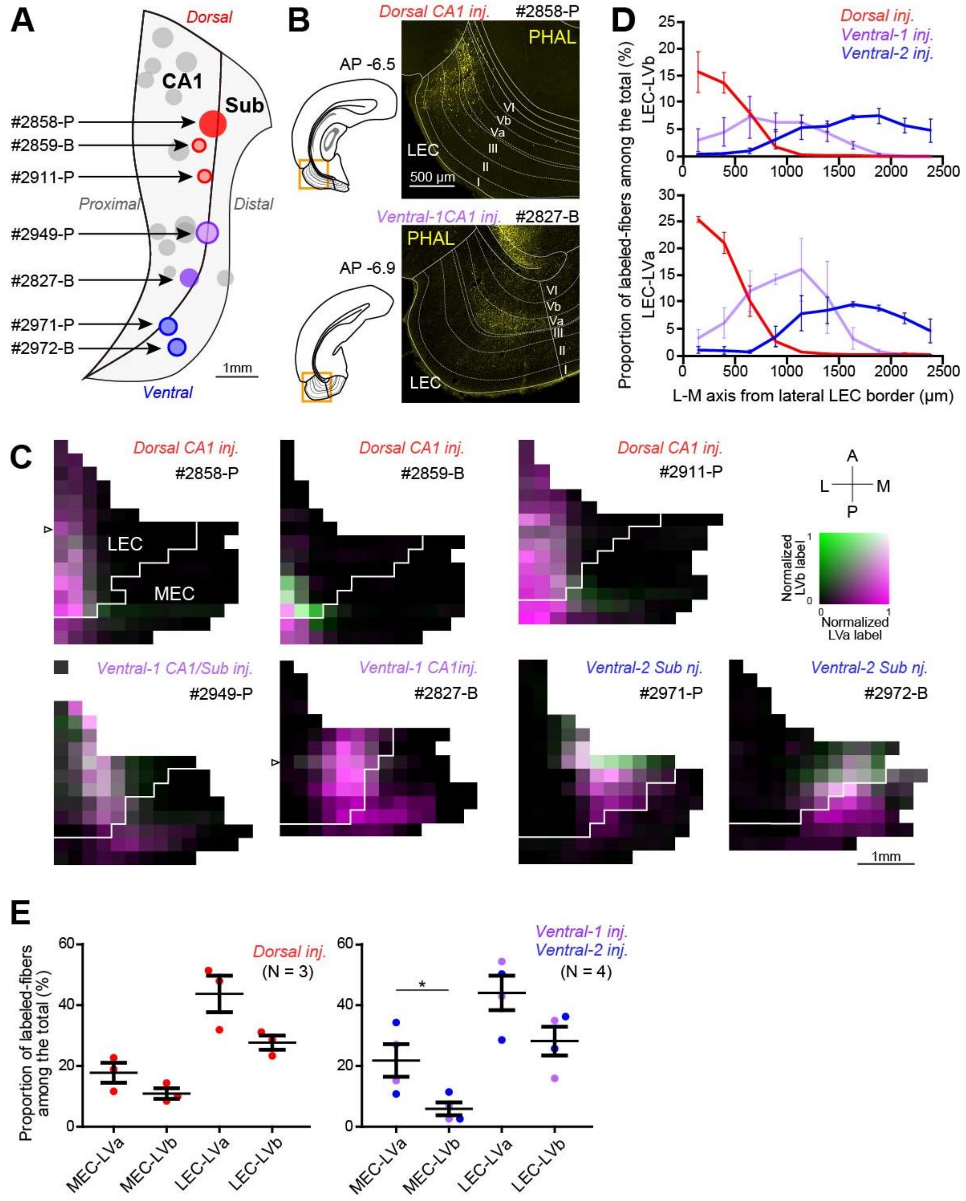


Figure S3. Topographical organization of hippocampal-LEC projections along the dorsoventral axis. (A) Two-dimensional unfolded map of CA1 and subiculum showing the positions of anterograde tracer (PHA-L or BDA) injection sites for rat samples in the coronal plane. Injection sites in the dorsal hippocampus are shown in red while injections in the ventral hippocampus are shown in purple (ventral-1) and blue (ventral-2). (B) Distribution of anterogradely labeled axons (yellow) in EC at different anterior-posterior (AP) levels in coronal sections. Images show samples injected in either dorsal (top, case #2858-P) or ventral CA1 (bottom, case #2827-B). (C) Seven representative two-dimensional density maps showing the patterns of anterogradely labeled axons in EC following anterograde tracer injections in either the dorsal or ventral hippocampus. The white arrowheads in #2858-P and #2827-B show the positions of images from B. (D) Proportion of labeled fibers in each subregion and sublayer among all labeled fibers in LVa and LVb along the dorsoventral axis of LEC. (E) Proportion of labeled fibers among all labeled fibers in MEC-LVa, MEC-LVb, LEC-LVa, and LEC-LVb for samples injected in the dorsal (error bars: mean ± standard errors; two-tailed paired t-test for MEC-LVa vs. MEC-LVb: $t2 = 3.66$, $p=0.15$, LEC-LVa vs. LEC-LVb: $t2 = 2.09$, $p=0.28$) and the ventral hippocampus (two-tailed paired t-test for MEC-LVa vs. MEC-LVb: $t3 = 3.56$, *$p=0.038$, LEC-LVa vs. LEC-LVb: $t3 = 2.01$, $p=0.43$).

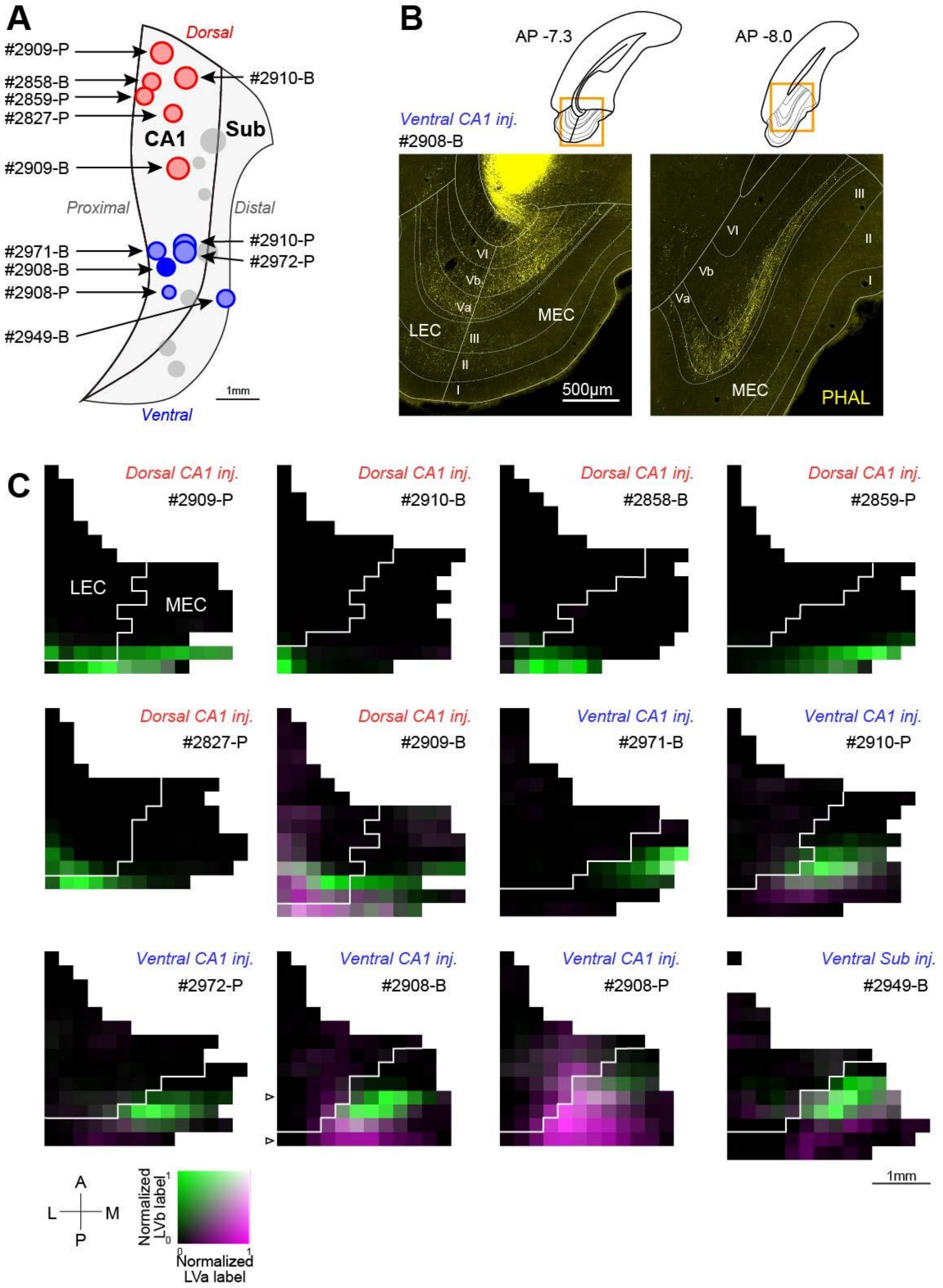


Figure S4. Hippocampal-entorhinal projections in the rat coronal plane originating from proximal CA1/distal subiculum. (A) Two-dimensional unfolded map of CA1 and subiculum showing the positions of anterograde tracer (PHA-L or BDA) injection sites for rat samples in the coronal plane. Injection sites in the dorsal and ventral hippocampus are shown in red and blue, respectively. (B) Distribution of anterogradely labeled axons (yellow) in EC at different AP levels in coronal sections after BDA injection in ventral CA1 (#2908-B). (C) Twelve representative two-dimensional density maps showing the patterns of anterogradely labeled axons in EC following anterograde tracer injections in either the dorsal or ventral hippocampus. The white arrowheads in #2908-B show the positions of images from B.

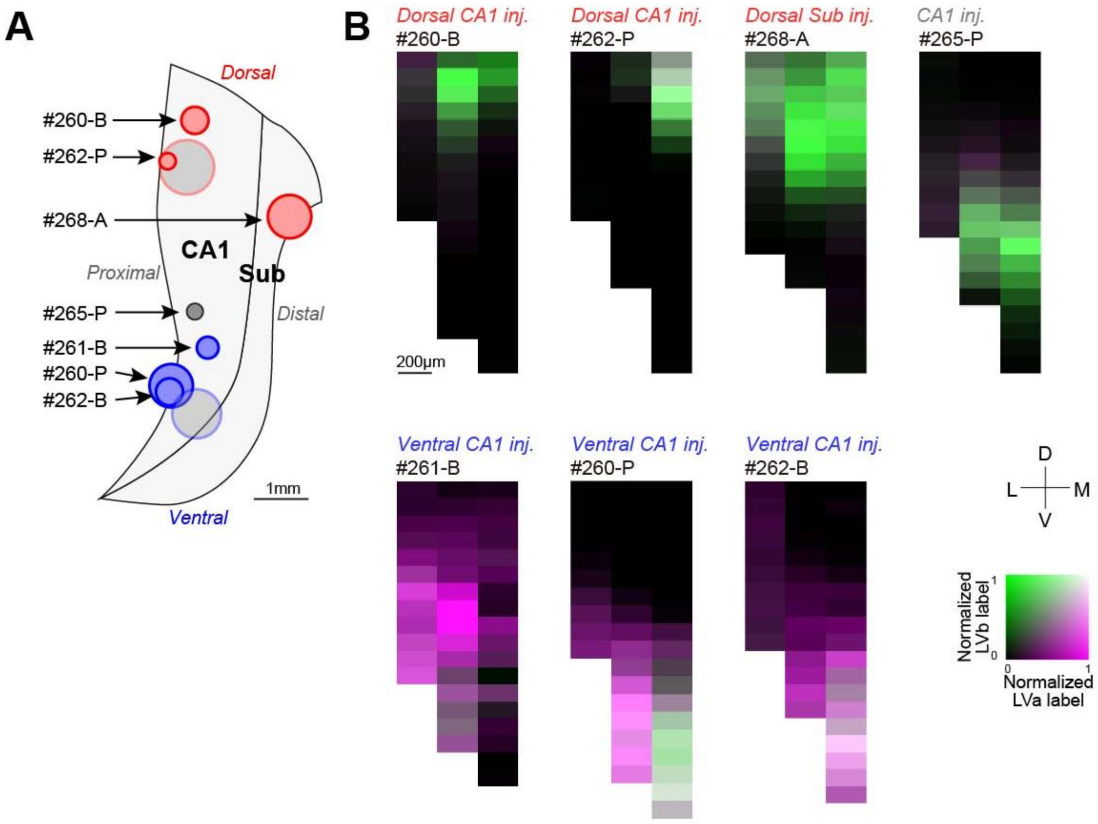


Figure S5. Related to Figure 1, shows label distribution patterns in unfolded EC maps for rat horizontal samples. #265-P, which was located in the intermediate hippocampus, was excluded from the analysis in Figure 2D.

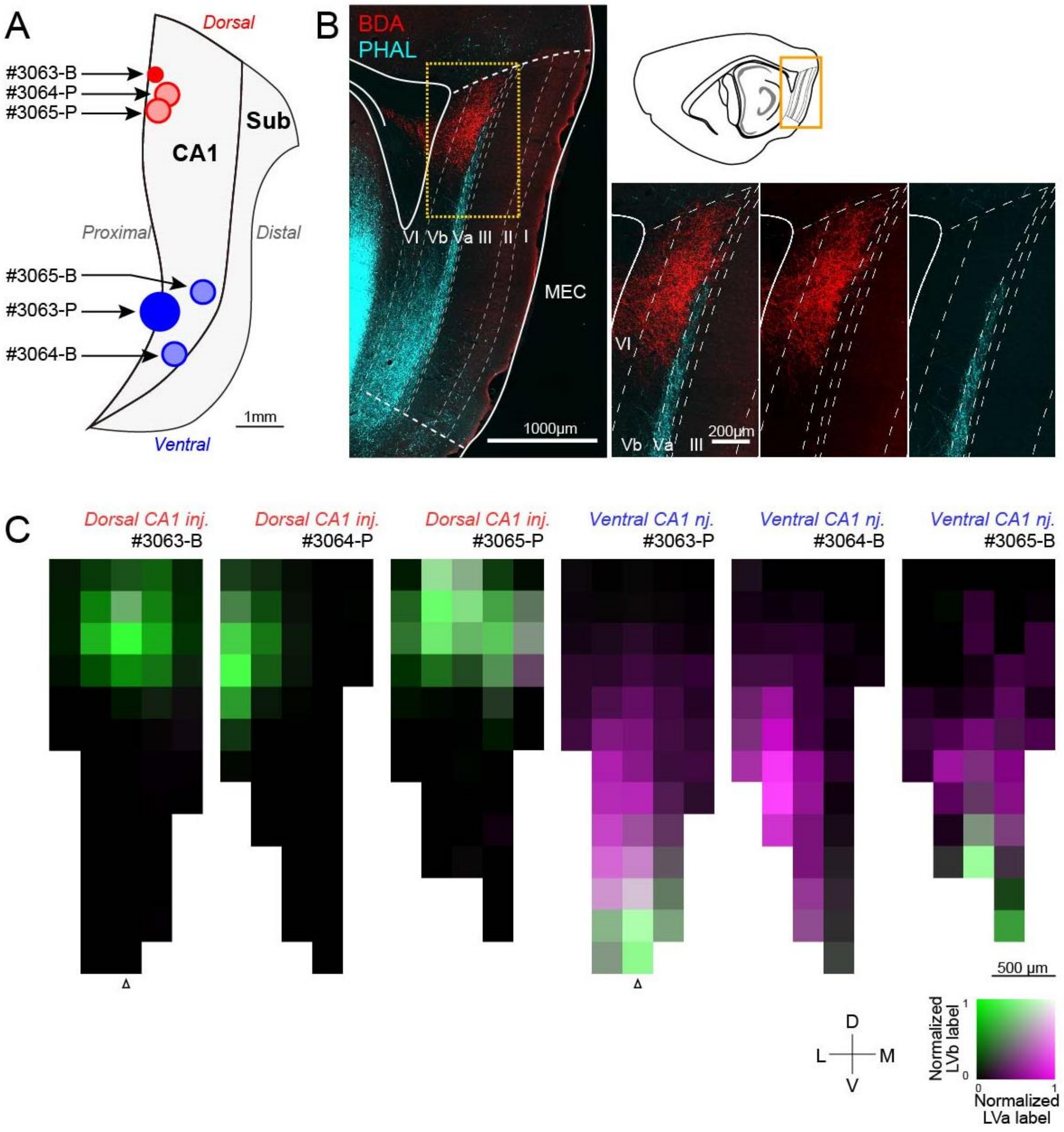


Figure S6. Hippocampal-entorhinal projections in the rat sagittal plane. (A) Two-dimensional unfolded map of CA1 and subiculum showing the positions of anterograde tracer (PHA-L or BDA) injection sites for rat samples in the sagittal plane. Injection sites in the dorsal and ventral hippocampus are shown in red and blue, respectively. (B) Fluorescent micrographs of MEC showing labeled axons originating from dorsal and ventral CA1 (red and cyan, #3063). (C) Six representative two-dimensional density maps showing the patterns of anterogradely labeled axons in MEC following anterograde tracer injections in either the dorsal or ventral hippocampus. The white arrowheads below the maps of #3063-B and #3063-P show the positions of images shown in B.

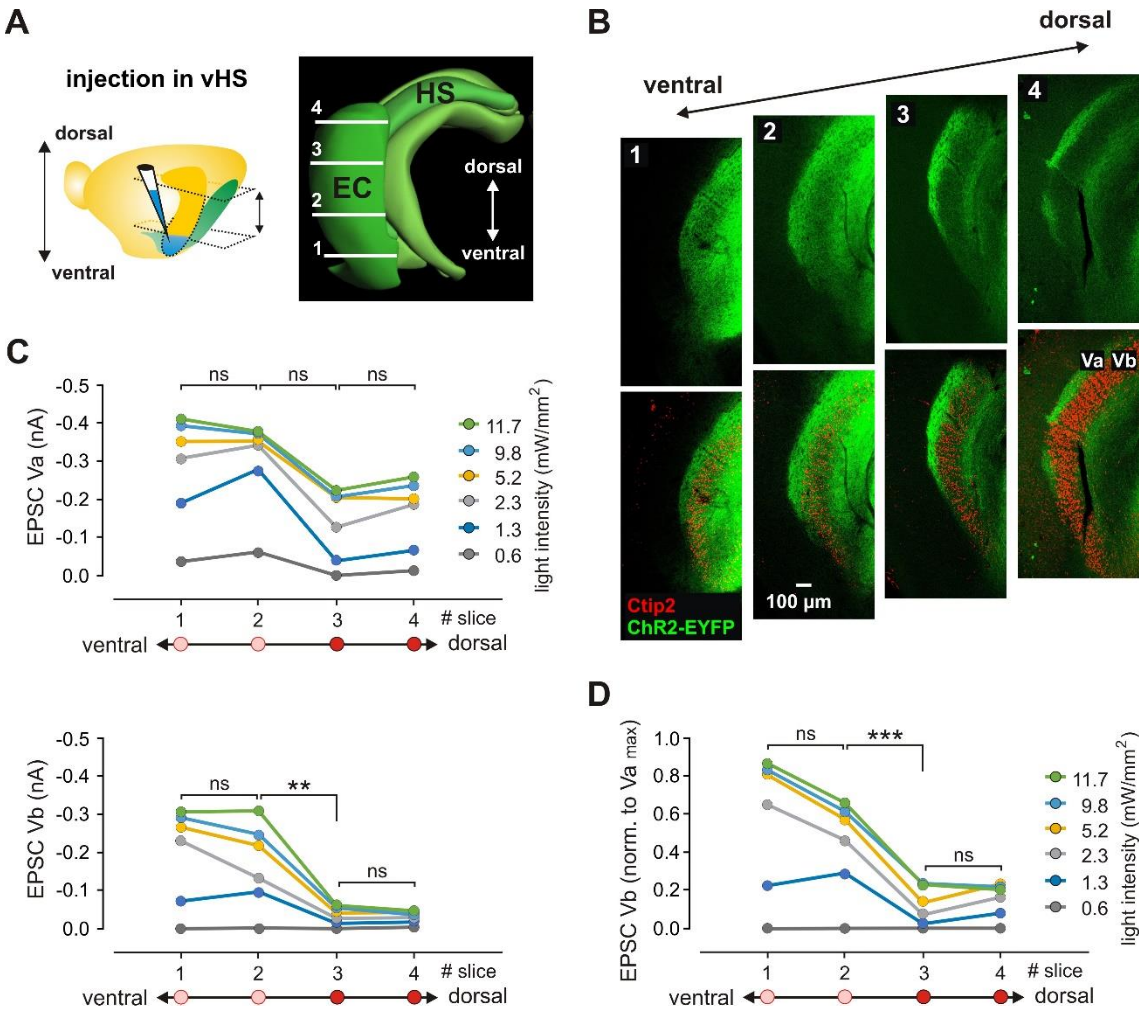


Figure S7. Functional connectivity between the ventral hippocampus and MEC LV along the dorsoventral axis. (A) Left: Illustration of the injection site (blue) in the ventral hippocampus (vHS). The approximate positions of horizontal sections used in experiments are indicated by arrows. Right: 3D-model of the mouse hippocampal formation (HS) and the adjacent entorhinal cortex (EC, posterior view). The image is modified from the Allen Brain Reference Atlas. The approximate locations and order of horizontal sections used in experiments are indicated with numbered horizontal lines. (B) Confocal images of a fluorescent staining of hippocampal axons expressing hChR2-EYFP at different levels of MEC. The bottom panels show images from the top overlaid with Ctip2 immunolabeling. Horizontal sections along the dorsoventral axis are indicated as shown in (A). Note the strong fluorescence of ventral hippocampal axonal fibers in Ctip2-negative LVa in the dorsal MEC (slice levels 3 and 4). (C) Quantification of synaptic responses of LVb neurons normalized to the highest LVa response at maximum light intensity (11.7 mW/mm$^2$) in each slice. Data are analyzed at four different section levels along the dorsoventral axis (1 to 4) for light pulses with increasing intensities. Note the strong differences in normalized LVb responses between the dorsal (levels 3 and 4), and ventral (levels 1 and 2) MEC. (D) Quantification of synaptic responses of LVb (left) and LVa (right) neurons recorded at four different slice levels along the dorsoventral axis. The panels show plots of EPSC amplitudes induced by light pulses with increasing intensities. All values are presented as median. Mann-Whitney test for response values at maximum light intensity: ***$p$<0.001; **$p$<0.01; ns, not significant.

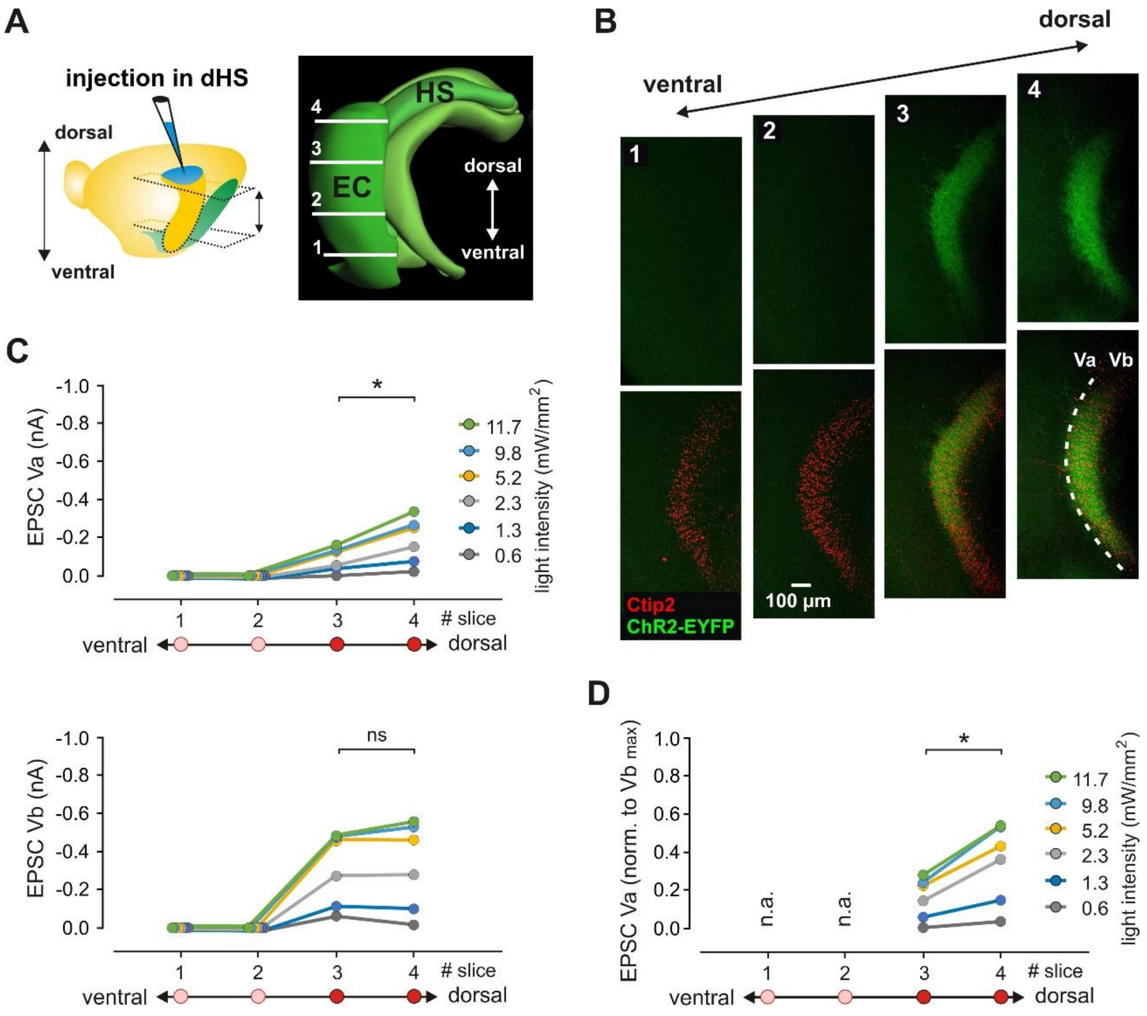


Figure S8. Functional connectivity between the dorsal hippocampus and MEC LV along the dorsoventral axis. (A) Left: Illustration of the injection site (blue) in the dorsal hippocampus (dHS). The approximate positions of horizontal sections used in experiments are indicated by arrows. Right: 3D-model of the mouse hippocampal formation (HS) and the adjacent entorhinal cortex (EC, posterior view). The image is modified from the Allen Brain Reference Atlas. The approximate locations and order of horizontal sections used in experiments are indicated with numbered horizontal lines. (B) Confocal images of a fluorescent staining of hippocampal axons expressing hChR2-EYFP at different levels of MEC. The bottom panels show images from the top overlaid with Ctip2 immunolabeling. Horizontal sections along the dorsoventral axis are indicated as shown in (A). Note the strong fluorescence of dorsal hippocampal axonal fibers in Ctip2-positive LVb in the dorsal MEC (slice levels 3 and 4) and the lack of fluorescence in the ventral MEC (slice levels 1 and 2). (C) Quantification of synaptic responses of LVa neurons normalized to the highest LVb response at maximum light intensity (11.7 mW/mm$^2$) in each slice. Data are analyzed at four different section levels along the dorsoventral axis for light pulses with increasing intensities. N.a.: not available (none of the neurons in ventral slices responded). (D) Quantification of synaptic responses of LVb (left) and LVa (right) neurons recorded at four different slice levels along the dorsoventral axis. The panels show plots of EPSC amplitudes induced by light pulses with increasing intensities. All values are presented as median. Mann-Whitney test for response values at maximum light intensity: *$p<0.05$; ns, not significant.
